# Intra-genomic genes-to-genes correlation enables bacterial genome representation

**DOI:** 10.1101/2024.06.12.598634

**Authors:** Jingjie Chen, Xuchuan Ma, Junwei He, Yingxia Wang, Yuji Ren, Li Qi, Liuyang Song, Lin Ruan, Cun Fan, Tao Huang, Jingbo Cheng, Xing Liu, Fang Chen, Yichen Huang, Haifeng Wang, Jiehui Chen, Yingying Pu, Na Zhao, Chunming Guo

**Affiliations:** Yunnan Key Laboratory of Cell Metabolism and Disease, Center for Life Sciences, School of Life Sciences, Yunnan University, Kunming 650500, China; Food Microbiology, Wageningen University & Research, Wageningen, the Netherlands; Deptartment of Biotechnology, Delft University of Technology, van der Maasweg 9, Delft 2629HZ, the Netherlands; The Department of Pathology, First affiliated Hospital of Kunming Medical University, Kuming, People’s Republic of China; The Department of Clinical Laboratory, First affiliated Hospital of Kunming Medical University, Kuming, People’s Republic of China; Department of Urology, Children’s Hospital of Chongqing Medical University, 136 Zhongshan Road, Chongqing 400014, PR China; Department of Urology, Shanghai Children’s Hospital, School of medicine, Shanghai Jiao Tong University; Key Laboratory in Software Engineering of Yunnan Province, Yunnan University, Kunming, People’s Republic of China; Big Data Research Center, University of Electronic Science and Technology of China, Chengdu, People’s Republic of China; Department of Urology, Yunnan Clinical Medical Center of Urological Disease, The Second Affiliated Hospital of Kunming Medical University, Kunming, China; State Key Laboratory of Cell Biology, CAS Center for Excellence in Molecular Cell Science, Shanghai Institute of Biochemistry and Cell Biology, Chinese Academy of Sciences, 320 Yue Yang Road, Shanghai, 200031, China; The State Key Laboratory Breeding Base of Basic Science of Stomatology & Key Laboratory of Oral Biomedicine Ministry of Education, School & Hospital of Stomatology, Medical Research Institute, Wuhan University, China

**Author notes:** These authors contributed equally (Jingjie Chen, Xuchuan Ma). Correspondence: Na Zhao < >, Chunming Guo < >.

**Keywords:** Pan-genome, Gene-Gene Correlation, Resistance Prediction

## Abstract

The bacterial pan-genome consists of core genes shared by all members of a taxonomy and accessory genes found in only a subset. The correlation among genes within the pan-genome could advance our understanding of evolution and tackle medical challenges. Here, we discovered a strong intra-genomic correlation among bacterial pan-genes within each of *Escherichia coli*, *Listeria monocytogenes*, *Staphylococcus aureus*, and *Campylobacter jejuni*. With a convolutional neural network assisted core genome knock-out simulation, we found that different combinations of fewer than 20 highly variable core genes could recover the sub-species type classified by complete core genome with accuracy >95%. This observation led us to test the genes-to-genes predictability: with more than 52,000 assemblies from each species, combinations of highly variable core genes could predict the sequence variants of other core genes (average accuracy >94%) within the same genome and could also predict sequence variants (average accuracy >91%) as well as the presence (average AUROC >0.91) of some accessory genes. Furthermore, combinations of highly variable core genes could also predict multiple antibiotic resistances (AUROC >0.80) in large published datasets of *E. coli*, *S. aureus*, and *Mycobacterium tuberculosis*. Collectively, we propose that genes within the same genome can strongly correlate with each other. Therefore, the strain phylogeny and the status of other genes could be uniformly represented by combinations of highly variable core genes, which could further represent certain phenotypes including *in vitro* resistance.

## Background

The bacterial pan-genome consists of core genes shared by all strains within a taxonomic unit and accessory genes found only in a subset. Core genes, characterized by their conservative presence, hold functions for basic survival, including but not limited to DNA metabolism, protein synthesis, and energy metabolism. Core genome is a well established concept proven experimentally using comparative proteomics; it defines bacterial lifestyle and the expression of core genes was relatively inflexible with respect to culture conditions^1^. Single-copy core genome based phylogeny is a frequently used method for strain subtyping, becoming increasingly achievable with the growing availability of assemblies and decreasing sequencing costs. Within-species single-copy core genome, containing hundreds to ore than thousands of genes, can classify strains into different core genome subtypes (cgSTs) at a high level of resolution while ensuring interlaboratory reproducibility^2,3^. The accessory genome, which includes genes that can be lost and acquired, facilitates the acquisition of functions. These genes include toxin genes, virulence factors (VFs), and antibiotic resistance genes (ARGs)^4–7^. The function acquisition partially explained the rapid adaptation of pathogens to new environments^8^. The accessory genome, as a gigantic pool of different functions, has been employed as input for machine learning based resistance prediction for certain antibiotics without a *priori* knowledge of mechanisms^9^.

Previous studies on relationship among genes^10–13^ had found that large amount of bacterial genes could display co-occurrence or avoidance patterns with one another in the accessory genome, and some studies on *Listeria monocytogenes* had presented a strong correlation between core genome based sublineage and accessory gene presence^14–17^. Beyond gene presence-absence patterns, internalin A (*InlA*) in *L. monocytogenes* is an example of correlation among SVs. Compared with original full-length version, truncated *InlA* is more specifically present in certain clonal complexes classified by seven housekeeping genes based Multi-Locus Sequence Typing (MLST)^14,15,18–22^. Further more, linkage disequilibrium (LD) as a classic study of gene correlation usually regards variant sites as a two-state (major/minor allele) object to carry out biallelic non-random association calculation. LD displayed strong distance dependency in both human and bacteria. In different species of bacteria, LD exhibited sharp initial decay within a few hundred base bps, but non-zero LD persists genome-wide due to bacterial clonal population structure^23–26^.

We noticed that studies exploring the relationships among genes in the pan-genome, particularly involving a large scale of genes and integrating correlation of both presence and sequence variants (SVs), were notably scarce. Herein, our study characterized the correlation among genes in the pan-genome by evaluating the genes-to-genes predictability. Genes are treated as a multi-state object whose status included their exact sequence and presence, were employed as input to infer the status of other genes in the same genome using convolutional nerual network (CNN) and also certain phenotypes such as *in vitro* antibiotics resistance. A large amount genes were involved, and high average predictability was achieved over the dataset of three bacterial phyla. Thus we believe this predictability is generalizable and will provide novel insights for pan-genome study.

## Results

### VFs and ARGs correlate with cgSTs in 137 *E. coli* strains

A correlation between VFs and ARGs and cgST were observed while studying local *E. coli* that caused Urinary Tract Infection (UTI). Ninety one strains were collected from patients in The First Affiliated Hospital of Kunming Medical College. Their assembled genomes were re-annotated together with 46 reference assemblies (Supplementary Table 1). A customized ortholog clustering pipeline OrthoSLC was employed to build gene clusters. For a given set of assemblies, gene clusters were categorized into accessory gene clusters (not shared by all assemblies) and core gene clusters (shared by all assemblies). If a given core gene cluster had one gene from each of the assemblies, the cluster is defined as ‘strict core’ gene cluster (StCGC); if any assembly had more than one gene in a core gene cluster, the core gene cluster is defined as ‘surplus core’ gene cluster (Supplementary Fig 1). The multi-sequence alignment (MSA) of 2170 StCGCs, were concatenated for phylogeny computation. Based on reference strains, the phylogeny was classified into 5 cgSTs with known references^3^ and 1 cgST without; this newly identified cgST was named as ‘F’ in our study (Figure 1a, Supplementary Data 1). All reference strains were in the same cgSTs as classified in the previous study, and *E. coli* collected from UTI patients were more frequently present in cgST B2 (53 out of 91)^3,27–31^. To interrogate whether B2 strains possessed specific VFs not carried by other cgSTs, the VFs of total 137 genomes were annotated, and more VFs of category adherence were detected in B2 strains (Supplementary Fig 2, Supplementary Data 1). This might indicate a better ability of urinary tract colonization of B2 strains which were more frequently isolated from UTI patients. Interestingly, some VFs were almost exclusively present only in certain cgSTs. For example, a fimbriae gene *cfaD*/*cfaE* was exclusively present in B1 strains (17/17 in B1, and 0/120 in non-B1 strains). Another group of genes, *shuY*, *shuA*, *shuS*, *chuT*, *chuX*, and *chuS* exhibited an absolute absence in cgST B1 and A (0/41 in B1 and A, 96/96 in other cgSTs) (Supplementary Table 2 for gene name and VFDB^32^ query correspondence).

**Fig 1:**
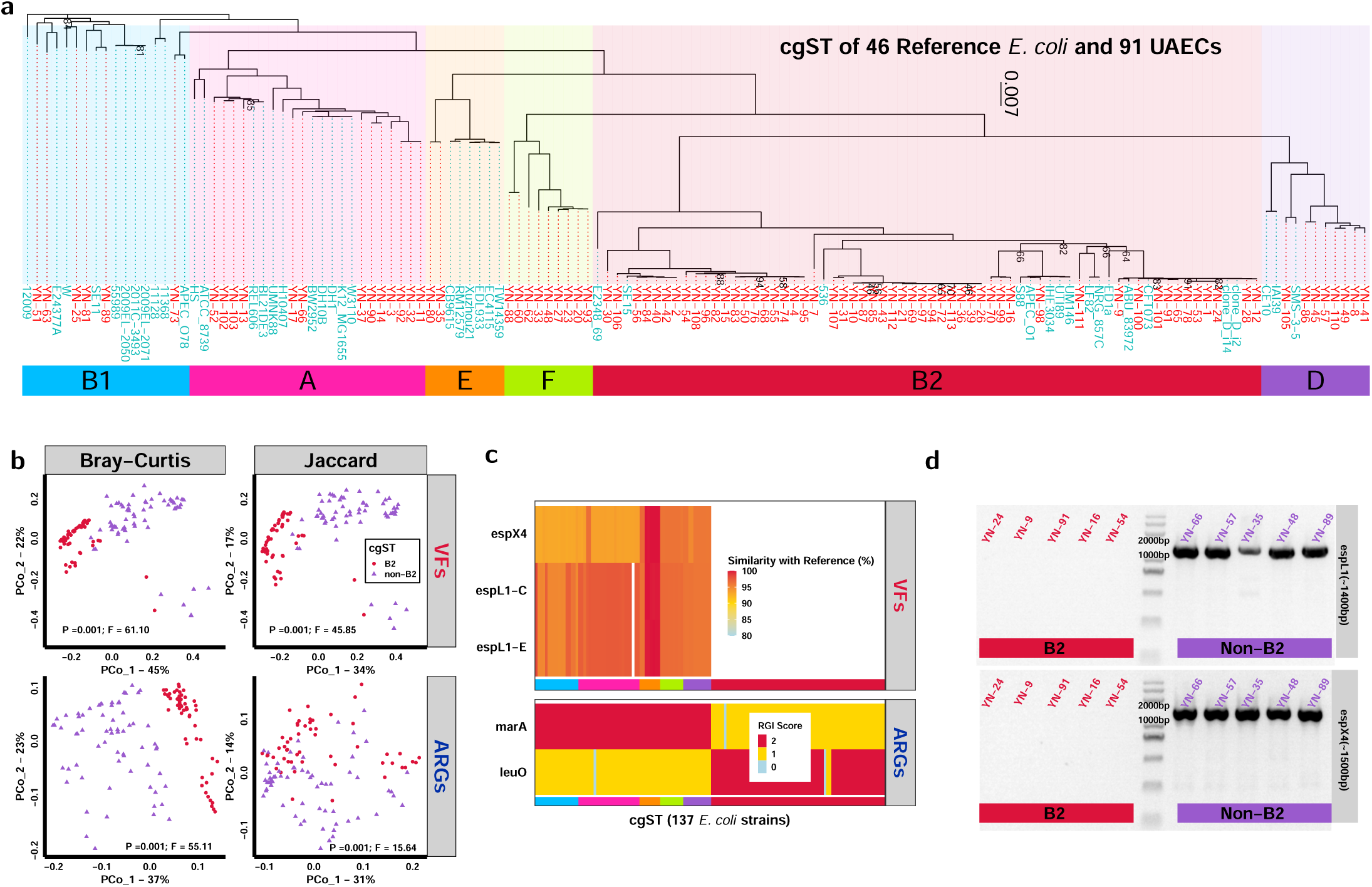
Both VF and ARG status correlate with cgSTs of UAEC. **(a) Single-copy core genome based phylogeny of 46 reference (cyan strain name) and 91 isolated (red name) *E. coli* strains.** The clade of each cgST was stained differently and corresponding cgST names are in the color bar below. Bootstrap values lower than 95 are shown. **(b) PCoA of VFs (first row) and ARGs (second row) using weighted (Bray-Curtis, first column) and unweighted (Jaccard, second column) algorithms.** The permanova based significance test between B2 (red dot) and non-B2 (purple triangle) strains is available on the bottom of each plot. **(c) Heatmap of genes specific to certain cgST selected by random re-trained LR models.** Color gradient in the VFs plot indicates the sequence similarity (<80% was regarded as absence of gene) between a corresponding gene of each strain and VFDB reference. *espL1-C* and *espL1-E* stand for *espL1* gene from *E. coli* CB9615 and EDL933 (Supplementary Table 2), respectively. RGI score of 0 stands for no resistance (not equivalent to gene absence); 1 stands for a strict hit, meaning potential resistance; and 2 stands for a perfect hit, indicating resistance. Scores were determined based on protein sequence similarity with reference and SNP site check by RGI^33^. The color on x axis stands for the cgST of each strain as in **a**. **(d) PCR based gene presence confirmation.** *espX4* had original length of 1581 bp in VFDB^32^ and its PCR target region is ∼1500 bp as indicated in sidebar; *espL1* original length is 1898 bp and target region is ∼1400 bp as indicated in sidebar.

To further examine the correlation between cgST and VFs, strains were divided into 2 major groups, cgST B2 (n = 68) and non-B2 (n = 69). Principal Coordinate Analysis (PCoA) was applied using genome VFs carriage status, including presence of VFs and SV of these present VFs which was reflected by similarity against reference sequences in VFDB^32^ (Figure 1b). The B2 and non-B2 groups exhibited a significant difference and a higher F value in Bray-curtis result than in Jaccard result, which suggested the cgST not only correlated with VFs presence but also correlated with sequence similarity against references. To find the VFs most significantly correlating with B2 or non-B2, the VFs status was used for B2 or non-B2 classification by Lasso Regression (LR) models, and VFs overlapping with StCGCs were excluded from input. The training was performed 50 times and each re-train started with randomly selected samples (70% from each cgST) as the training set. Test set AUROC of 1.00 was achieved in all 50 times of re-trains. By extracting the average weights that LR models gave to each VF over 50 re-trains, all VFs with an average weight of more than zero were displayed in Figure 1c. The result showed that *espX4* as a Type III secretion system (TTSS) secreted effector, exhibited exclusive (69/69) presence in non-B2 cgST. Similarly, two homologs of TTSS secreted effector *espL1* from *E. coli* CB9615 and EDL933 (Supplementary Table 2) showed almost exclusive (68/69) presence in non-B2 cgST. By targeted PCR (Figure 1d) and Sanger sequencing, those bands with correct length present in non-B2 genomes turned out to be corresponding sequences. These findings indicated a correlation between VF presence and cgST.

Besides VFs, ARGs were annotated using Resistance Gene Identifiers (RGI) and CARD database^33^ (Supplementary Data 1). The PCoA on ARGs status showed a significant difference between B2 and non-B2, and a higher F value in Bray-curtis result than in Jaccard result could suggest that RGI score as an indirect reflection of SV of ARGs is correlated with cgST (Figure 1b). Using ARGs status as input, the re-train based cross validation in predicting cgST of each strain returned two genes *marA* and *leuO* (Figure 1c). Both genes were present in almost all analyzed strains (136/137 for *marA* and 135/137 for *leuO*), and the RGI score of these two genes strongly correlated with cgST. *marA* was only scored 2 (68/68) in non-B2 strains and *leuO* was almost exclusively scored 2 in B2 strains (66/69). Hence, the SVs of some ARGs were specifically associated to cgST with a pronounced manner.

### Core-to-accessory genome correlation is present in multiple phyla

To confirm whether the correlation is present in more cgSTs and is ubiquitous in bacteria kingdom, we included 4 species, *E. coli*, *L. monocytogenes*, *S. aureus*, and *C. jejuni*. The four species were selected based on diversity and assembly availability consideration. *L. monocytogenes* and *S. aureus* are Gram positive while *E. coli* and *C. jejuni* are Gram negative. They originated from three Phyla: *Pseudomonadota*, *Bacillota*, and *Campylobacterota*. And these two *Bacillota* species *L. monocytogenes* and *S. aureus* are from different families, *Listeriaceae* and *Staphylococcaceae*, respectively. 65,840 *E. coli*, 52,090 *L. monocytogenes*, 54,817 *S. aureus* and 54,905 *C. jejuni* assemblies (Supplementary Data 2) were included in the analysis as Species Assembly Collection (SAC). To obtain sufficient assemblies and better represent more cgSTs, we adopted a strategy of pseudo-labeling based dataset construction (Figure 2a). In the workflow, Dataset I consists of 600 to 900 assemblies which were randomly selected from the SAC for each species, and a random forest (RF) model trained with Dataset I (Supplementary Data 3) was employed to predict the cgST as pseudo-labels for all members in SAC inputting with VF status of assemblies (Supplementary Data 2). The generated pseudo-labels were utilized to assign 200 assemblies to each cgST of Dataset II (Supplementary Data 3), including some cgST references selected from Dataset I. The result of Dataset II construction is shown in Figure 2b for *E. coli* and Supplementary Fig 3-5 for the other 3 species. The identification of cgST in Dataset II were referenced by strains selected from Dataset I (Supplementary Table 3-6). In Dataset II, 478 StCGCs for *E. coli*, 1,082 for *L. monocytogenes*, 672 for *S. aureus* and 522 for *C. jejuni* were constructed (Supplementary Data 7). The assignment of genomes from SAC to cgSTs showed a high accuracy. Besides cgST A and B in *S. aureus*, all other cgSTs among species of Dataset II were assigned 200 ± 5 assemblies. Therefore, these results indicate a correlation between cgST and VF status.

**Fig 2:**
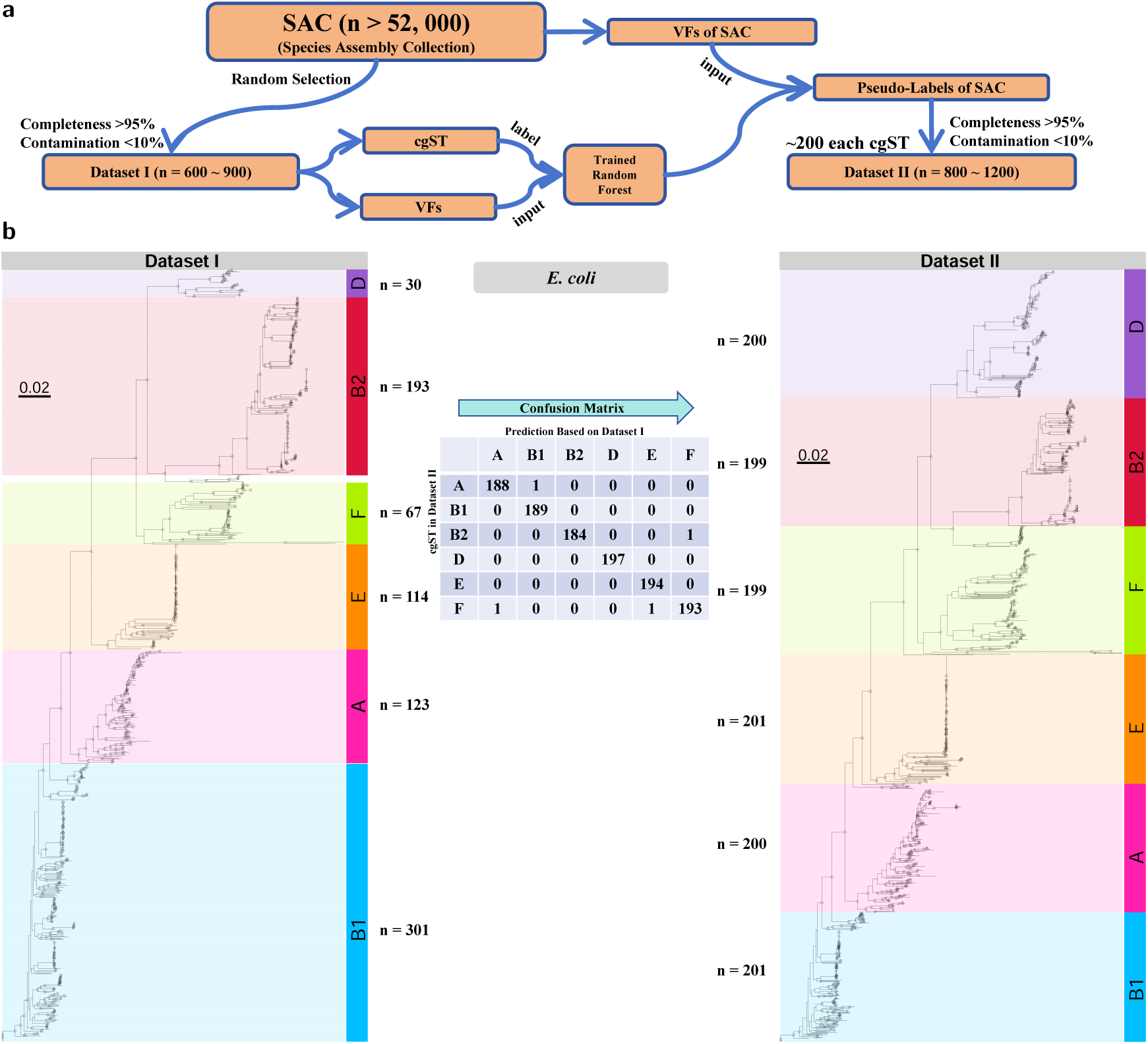
Accurate prediction of cgST by accessory gene status across three phyla. **(a) The schematic workflow of Dataset construction.** Pseudo-labels for all assemblies in SAC were generated by the random forest model trained using Dataset I. These pseudo-labels were used to assign ∼200 assemblies to Dataset II (including reference with known cgST from Dataset I). **(b) cgST result of Dataset I (left) and II (right) of *E. coli*.** cgSTs of clades were identified using reference strains (Supplementary Table 3-6). The uncolored branches in Dataset I were not classified into any cgST and not used in RF model training. The dashed branches were of large length and were manually shortened to help visualization. The nodes with bootstrap value 95 are marked with circles and those with bootstrap value >80 and <95 were marked with triangles. The tables in the middle of **(b)** describes the confusion matrix which excludes the reference strains (from Dataset I) and amonut of strains next to each cgST of Dataset I and II. Results for *L. monocytogenes*, *C. jejuni*, *S. aureus* are available in Supplementary Fig 3-5.

With sufficient assemblies in each cgST of Dataset II (Supplementary Data 7), the accessory genes that most significantly correlate with certain cgSTs was analyzed. By using VF or ARG status (Supplementary Data 3 and 4) as input, 50 retrains were applied on binary classification of each cgST using LR models. Notably, genes overlapping with StCGCs were excluded from prediction input. Genes with high average weights in trained models were displayed in Supplementary Fig 6. The average weights of all VFs or ARGs could be seen in Supplementary Data 5. The results in Supplementary Fig 6 suggested that cgST could be predicted using VF status with an average AUROC higher than 0.96 for all cgSTs and status of many accessory genes apparently correlated with cgST. Relatively low average AUROC (<0.90) when using ARGs as input in *L. monocytogenes* and *C. jejuni* could result from the lack of known ARGs in these species for cgST prediction. To date, *E. coli* had >250,000 total assemblies in NCBI database, and there were in total 643 VFs, 249 ARGs in Dataset II. In contrast, there were less than 100,000 total assemblies in NCBI assembly database for each of *L. monocytogenes*, *C. jejuni*, and *S. aureus*, and only 50, 151, and 122 VFs as well as 22, 37, and 79 ARGs were found in Dataset II, respectively. Despite these variations, correlation among cgST, VFs, and ARGs could be observed across Phyla and in more cgSTs.

### Core gene subsets recover the phylogeny with CNN

According to previous studies^10–13^, certain bacterial genes had significant co-occurrence/avoidance patterns. Thus, the correlation between accessory gene status and cgST in Supplementary Fig 6 might actually result from the gene co-occurrence phenomenon. Because some StCGCs might actually be a misclustering result of two or more accessory gene clusters which had distinct functions (heterofunctional cluster). If a heterofunctional cluster is decisive for strain cgST classification, the cgST specific accessory gene presence could with another gene (Supplementary Fig 7). On the other hand, if a certain cgST was decided by one or multiple StCGCs and each consisted of genes with a same function (homofunctional cluster), this cgST specific accessory gene presence shall not be gene co-occurrence but rather a correlation between core gene SV and accessory gene presence (Supplementary Fig 7).

To verify if observed correlation is gene co-occurrence or not, we delved into all StCGCs and attempted to find a subset of StCGCs, which were homofunctional and could still recover the complete core genome cgST. This was confronted with two challenges, the large number of subset StCGC combinations and extensive computation time of phylogeny with manual cgST identification. To overcome these obstacles, a simple Cluster Variation Level (CVL) algorithm was designed to value the intra-cluster diversity and infer the potential that each cluster could decide the cgST classification (Figure 3a). A 5-layer CNN as model A (Supplementary Fig 8), which could be reproducibly trained to generate high test set performance (accuracy >98%) with complete core genome as input, was employed. We carried out cross validation based cgST recovery evaluation by inputting different combinations of StCGCs. Within each species, the samples of Dataset II were separated into three equal size sub-groups. Thus, the cross validation was performed three-folded with three re-trains with Xavier random initialization^34^ on each fold, meaning a total of nine re-trains to evaluate each input StCGC combination. To input different subsets of StCGCs, the complete core genome dimension was maintained but positions of non-input genes were all set to zero, as knock-out (KO) simulation. In Supplementary Fig 9 first row and Figure 3b first row, by increasing the number of StCGCs as prediction input starting from the highest CVL rank, the test set performance was compared with when complete core genome and all-zero matrix were input. We found that 6 highest CVL StCGCs for *E. coli*, 11 for *L. monocytogenes*, 10 for *C. jejuni* and 5 for *S. aureus* were needed to recover cgST based on complete core genome with test set accuracy >98% (Supplementary Data 8), and these top CVL StCGCs are defined as hub StCGCs. This outcome corresponded to the phylogenetic tree of *E. coli* constructed using hub StCGCs only (Supplementary Fig 10), in which the cgST result were accurately recovered (1197/1200) comparing with the phylogeny structure based on complete core genome. After function annotation with eggnogmapper^36^, all these hub StCGCs turned out to be of homofunctional on protein sequence level (Supplementary Data 7). This result could primarily illustrate that cgST in Dataset II could be decided by homofunctional StCGCs. Therefore, the correlation that accessory gene status was specific to certain cgSTs did not result from gene co-occurrence.

**Fig 3:**
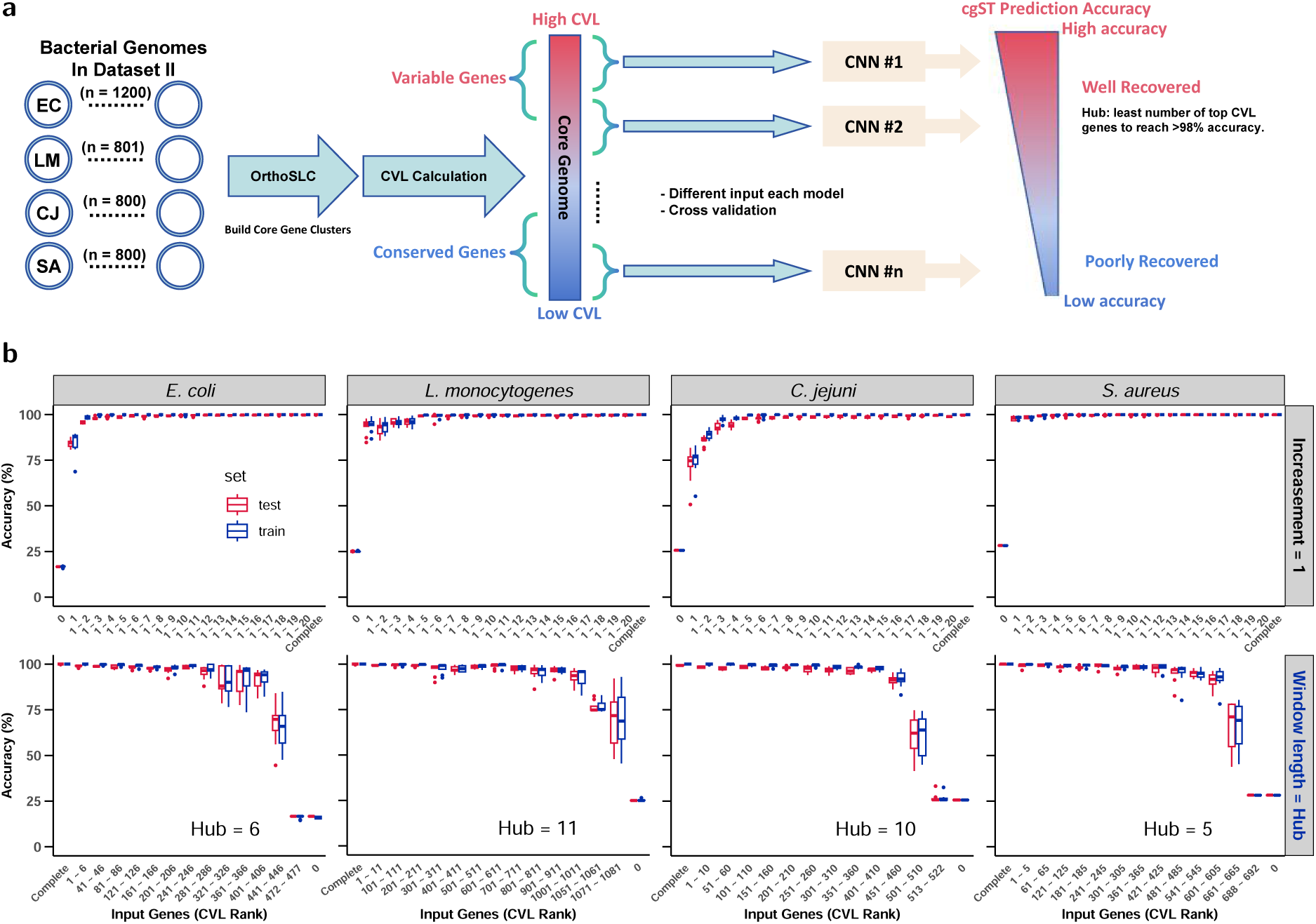
Limited core genes can accurately recover cgSTs with CNN in cross validations. **(a) schematic workflow:** re-annotated genomes of Dataset II (EC: *E. coli*, LM: *L. monocytogenes*, CJ: *C. jejuni*, SA: *S. aureus*) were sent to construct gene clusters where StCGCs were ranked by their CVL. The core genome was concatenated from high CVL StCGCs (genes more variable among strains) to low CVL gene clusters (genes more conserved among strains). Different KO combinations allowed the test of the number of hub genes (least top CVL genes needed to recover complete core genome cgST with >98% accuracy) as in **b** first row and Supplementary Fig 7 first row. By using different windows of available core genes in prediction (**b** second row), the ability to recover complete core genome cgST was tested on a wider range of core genome. Each box of distribution represents the performance of nine re-trains (three-fold, three random initialization each fold) of train set (red) and test set (blue). The CNN model adopted model A as in Supplementary Fig 7.

To investigate if homofunctional StCGC based cgST recovery could also be achieved with non-hub StCGCs, their function annotation was performed: only two StCGCs in *E. coli* (the 106^th^ and 134^th^ in CVL rank) and one in *C. jejuni* (ranked 163^rd^) were heterofunctional among the four species (Supplementary Data 7). This result suggested that most StCGCs were homofunctional, so the same cross validations same as Figure 3b first row was applied to evaluate more combinations of input (one StCGC from each of *E. coli* and *L. monocytogenes* with 0 CVL value was excluded from input). In Figure 3b second row (Supplementary Data 8), *E. coli* result showed that 95.48±3.58% cross validated average accuracy of test set could still be achieved inputting with 281^st^ to 286^th^ highest CVL StCGCs. These 286 highest CVL StCGCs represented 59.96% of all *E. coli* StCGCs (least among four species), so the hub StCGCs were not the only options for recovering the cgST of the complete core genome. In Supplementary Fig 9 second row, when the window length of input was increased to 20 genes, better performance could be achieved, the test set performance of 401^st^ to 420^th^ highest CVL StCGCs reached 97.33±0.99% for *E. coli*. In Supplementary Fig 9 third row, around 100, 250, 150, 200 lowest CVL rank StCGCs for *E. coli*, *L. monocytogenes*, *C. jejuni* and *S. aureus*, was required to recover complete core gene cgST with accuracy >98% (test set). Performance of these input windows corresponds to Supplementary Fig 10-13 of *E. coli* phylogeny structures. When top StCGCs were used for phylogeny construction, the cgST result for all strains was recovered almost perfectly although the detail was not identical to the complete core genome phylogeny. When lower-ranked StCGCs were input, only certain cgSTs were still recovered well. In Supplementary Fig 13, when 100 lowest CVL *E. coli* StCGCs were input, 1192 out of 1200 were correctly assigned. These results suggested that complete core genome cgST could be recovered using many combinations of StCGCs while these StCGCs are homofunctional. Thus, cgST specific SV and presence of an accessory gene did not result from gene co-occurrence.

### Core gene subsets predict other genes in the same genome

In Supplementary Fig 6, the observation of correlation between cgST and accessory gene status was limited because the cgST was only divided into several general classes. To observe the correlation between core and accessory gene status more directly, accessory gene presence prediction was attempted using StCGCs as input and CNN model B from Supplementary Fig 8. Within each species, genes present in >5,000 and absent in >5,000 SAC assemblies per species were evaluated for presence predictability. Two-fold cross validations were performed with two randomly initialized re-trains on each fold, so the test set performance of four re-trains was employed to evaluate the predictability of a single gene. To balance the computational workload and sample size, the input dimension was reduced. A top window in CVL rank list with only hub StCGCs were employed as input and two other windows were selected from the middle and lower part of CVL rank list (Figure 4a). Middle and lower windows included more genes to reach a similar sequence length of the top window. These three windows included a KO version and a complete version of input. In addition, a window using housekeeping genes was included (Supplementary Table 8).

**Fig 4:**
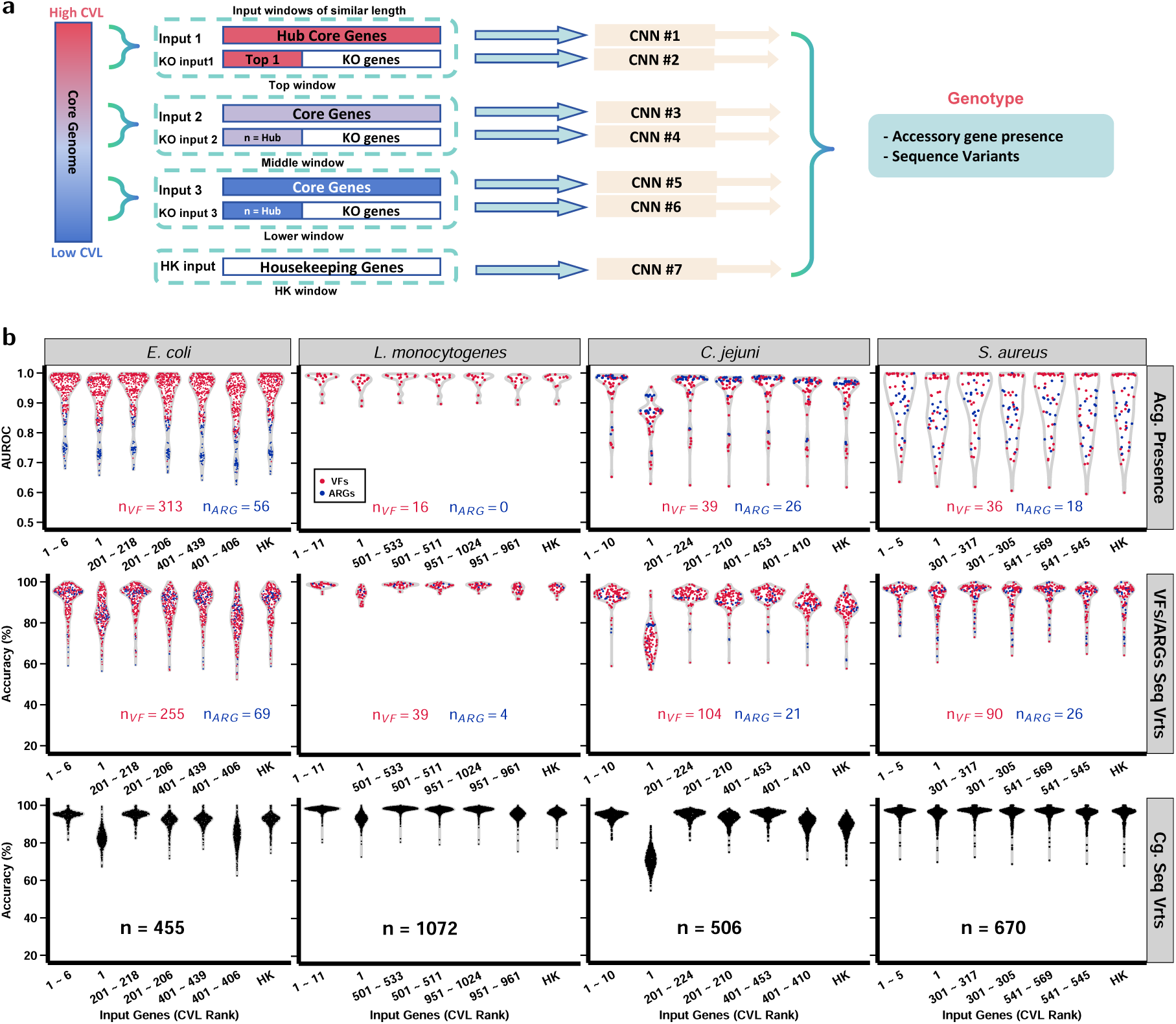
Limited core genes can predict status of other genes within the same genome with a high average accuracy. The workflow (**a**) included windows of similar length as input, a top window with all hub core genes each species and 2 windows using middle and lower end CVL rank StCGCs. All windows included a KO version and a complete version of input. KO input for top windows kept only the highest CVL StCGC as input; middle and lower windows kept only first (amount of hub) genes. In addition, a window of housekeeping genes (Supplementary Table 8). Each point represented the average test set performance (two-fold, two random initialization) of four re-trains for accessory gene (Acg.) presence prediction (**b** first row) and SV (Seq Vrts) prediction of VFs/ARGs as well as core genes (Cg.) (**b** second, third row). The number of genes being predicted are available in each plot. For each gene included in presence prediction (**b** first row), there should be >5,000 samples from SAC with the gene present in the assemblies while assemblies without the gene present should also be >5,000. For each gene included in SV prediction (**b** second, third row), there should be at least two SVs owned by >2,000 assemblies in SAC. The CNN model adopted model B in Supplementary Fig 8.

Among the four species, all complete version of top windows reached average test set AUROC >0.91 (0.91±0.09 in *S. aureus* as the lowest of four species) in predicting accessory gene presence (Figure 4b first row, Supplementary Data 9). Notably, presence of accessory genes could not always be accurately predicted. For example, ARGs in *E. coli* had an average AUROC 0.79±0.06 using top window as input. Apart from complete top windows, similar performance could be achieved with complete middle, lower and housekeeping windows. This suggested that the presence of accessory genes could be inferred by different combinations of StCGCs. Besides the KO input of the top window in *C. jejuni*, prediction, KO input of all windows exhibited a slightly lower performance compared with their complete versions. This suggested that inferring the presence of some accessory genes required a limited amount of core genes which could be selected from a large range.

Apart from presence, SV prediction of VFs and ARGs was also tested. Any gene to be predicted had at least 2 prevalent SVs that were present in more than 2,000 assemblies of the SAC. Assemblies were grouped into classes owning different prevalent SVs of the gene to be predicted, and a class ‘other’ of assemblies holding all other non-prevalent SVs. A two-fold cross validation was performed, and each fold had two re-trains with random initialization, so a total of 4 re-trains for evaluation. In Figure 4b second row (Supplementary Data 10), all complete top windows reached average accuracy >91% (91.98±6.91% in *E. coli* as lowest among four species) and similar accuracy in other complete windows. Performance of KO input showed some decrease comparing with their complete versions. For *E. coli*, performance of top window input reached 96.90±5.85% when the class ‘other’ was excluded from the prediction (Supplementary Fig 14 first column). These results exhibited that there is a correlation between the core gene status and the status of accessory genes from the same genome, this correlation could be reflected by both accessory gene presence and SV prediction by input of combinations of limited amount of StCGCs from a large range.

Besides the correlation between core and accessory genes, the observation in Figure 3b second row that different StCGC combinations could predict the same cgST led to a hypothesis: core genes within a same genome could predict the SV of each other. To test this hypothesis, the same input of StCGC combinations in Figure 4b first and second row were employed. Notably, if the gene to be predicted is one of the input genes, the corresponding positions of the input would be set as zero. Self prediction was skipped when first gene in CVL rank was the only input and the gene to be predicted at the same time. In Figure 4b third row (Supplementary Data 10), all top windows showed an average accuracy >94% (94.47±2.35% in *C. jejuni* as lowest among four species), and similar accuracy could be achieved using other complete windows as input. For *E. coli*, the top window result increased from 94.65±2.40% to 97.61±2.19% when class ‘other’ was excluded (Supplementary Fig 14 first column). Notably, some prevalent SVs could have only one nucleotide difference while the prediction still reached high accuracy (Supplementary Fig 15, 16). These results suggested that correlation among genes was not limited between core and accessory genes, different combinations of StCGCs could infer other genes with a high accuracy.

Notably, this predictability involved a large range of genes in the pan-genome. In *E. coli* SAC dataset. We generated DNA sequence embeddings using a pre-trained model B (Supplementary Fig 8). By passing each sequence through the CNN and extracting activations from the penultimate layer, we obtained compact 688-dimensional feature vectors. These embeddings were then used as fixed inputs to the final fully connected layer, allowing us to retrain only this layer during fine-tuning. Genes in *E. coli* dataset II with presence rate <100% were extracted from SAC, and status of these pan-genes were predicted with embedded input. Among these pan-genes, presence prediction of 3605 accessory genes with AUROC 0.91±0.09 and SV prediction of 4913 pan-genes with accuracy of 91.39±7.17% (Supplementary Fig 14 second column, Supplementary Data 11). Here, we adopt the term genes-to-genes (g2g) correlation to conceptualize the observation. The g2g correlation means that the status of a given gene, including presence and SVs, correlates with the status of other genes from the same genome. Such correlation can explain the phenomenon that the status of some genes can be accurately predicted by other genes from the same genome.

### Core gene subsets predict antibiotic resistance

Core genes exhibited great power in inferring the status of other genes, implying a potential to represent a genotype closer to whole genome with limited parts and to better encode phenotypic traits. To test this potential, published datasets^9,37–40^ (Supplementary Table 6) with clear antibiotics resistance labels (Supplementary Data 12) were selected from BV-BRC^41,42^ to obtain assemblies of *E. coli*, *S. aureus* and *Mycobacterium tuberculosis*. We emplyed the same three-fold three random initialization cross validation as in Figure 4, and observed deviation among re-trains correlating with low sample size, a larger sample size showed more consistency over re-trains (Supplementary Fig 17a). However, the large sample size did not necessarily lead to high performance (Supplementary Fig 17b). For example, Amoxicillin-Clavulanate (AMC) and Ampicillin (AMP) in *E. coli* of published datasets had average AUROC lower than 0.80 while there were at least 339 assemblies representing susceptible and resistant strains (). Interestingly, predictions of resistance to multiple drugs reached average AUROC >0.80 in at least two input windows, and the prediction of some resistance exhibited AUROC >0.90, including Ciprofloxacin (CIP) and Methicillin (MET). Particularly, CIP resistance exhibited prediction AUROC >0.90 across three species based on the data from three different studies^9,38,40^.

For AMP resistance prediction in *E. coli*, AUROC 0.70±0.01 for dataset of Davies et al.^37^ and 0.73±0.02 for dataset of Moradigaravand et al.^9^ were achieved inputting with hub StCGCs (Figure 5b, Supplementary Data 13). However, Davies et al.^37^ achieved AMP prediction sensitivity 98% by simply predicting strains owning beta-lactamase genes (e.g., *OXA-1* and *TEM-30*) as AMP resistant. Therefore, the presence prediction performance of *TEM-30* (AUROC 0.76±0.01, Supplementary Data 9) corresponded to the low AMP resistance performance when inputting with hub StCGCs. In contrast, the high performance (AUROC >0.90) on CIP and MET resistance prediction in *S. aureus* and *E. coli* using hub StCGCs corresponded to accurate prediction on key genes. The MET resistance related genes *mecR1, mecI*, *mecA* available in our study (Supplementary Data 9) had presence prediction AUROC of 0.97±0.01, 0.97±0.01 and 0.87±0.01, respectively, inputting with hub StCGCs. Sequence variant prediction accuracy of *gyrA* (Supplementary Data 10) reached 0.80±0.01 in *S. aureus* and 0.95±0.01 in *E. coli* (inputting with hub StCGCs), which corresponded to CIP resistance prediction performance. Notably, the performance of *M. tuberculosis* Isoniazid (INH) and Rifampicin (RIF) resistance prediction using housekeeping gene input was higher than other input windows. Since the most resistance-related genes for INH and RIF, *katG* and *rpoB* are included in the housekeeping genes, respectively, this accurate resistance prediction should not result from a better ability of housekeeping genes in inferring the status of other genes than other input windows. These results suggested that while phenotype prediction is still challenging in general, but certain antibiotics resistance could be well inferred implying the correspondence between phenotype and genotype which was represented by limited core genes.

**Fig 5:**
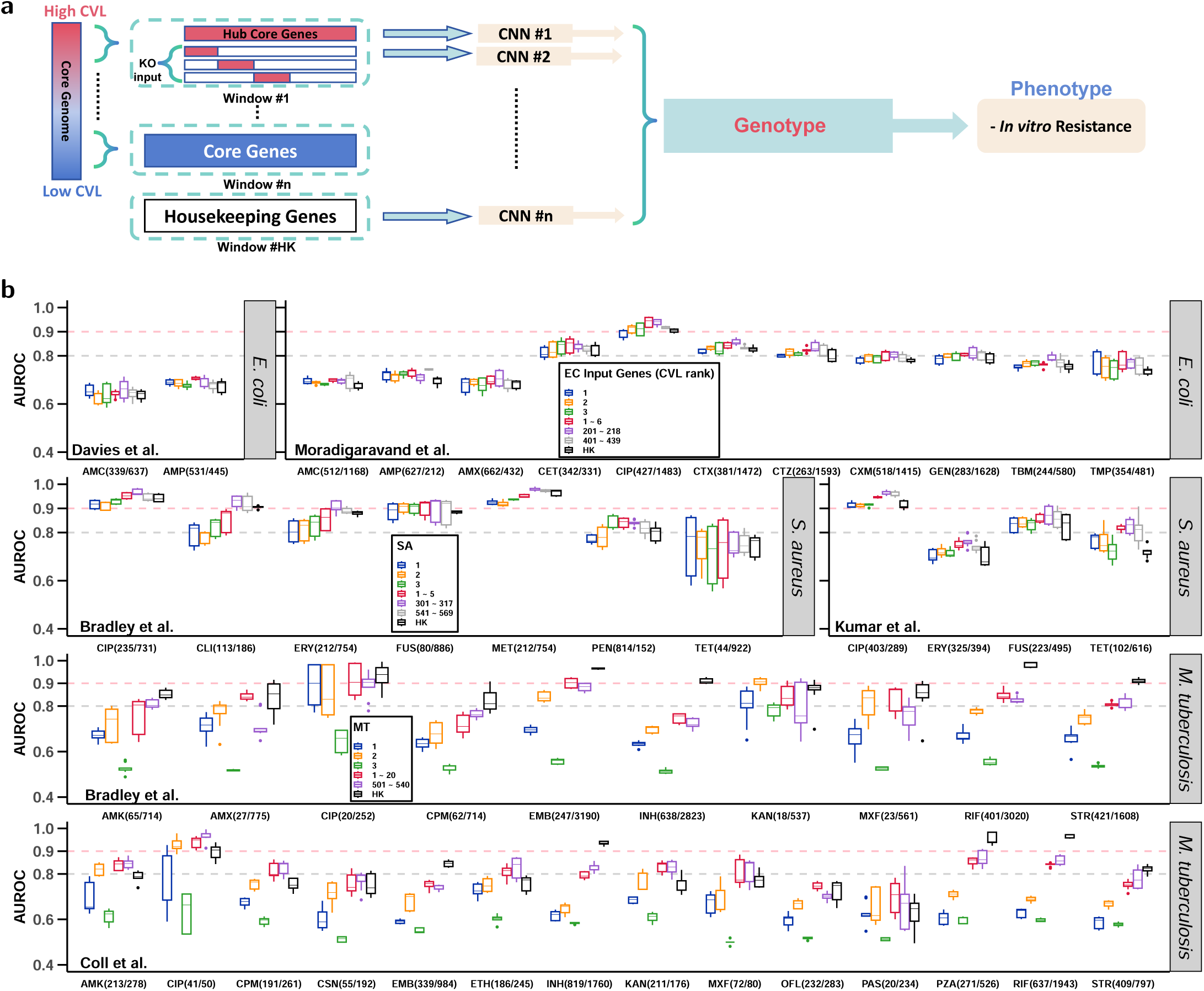
Limited core genes predict antibiotics resistance. The input windows shown as in **(a)**, included a complete top window of hub core genes and three KO input with three highest CVL genes kept, respectively. Both *E. coli* (EC) and *S. aureus* (SA) included 3 windows as in Figure 4, while *M. tuberculosis* (MT) includes only 2 windows. **(b)** Results of each species included a input window of housekeeping genes. Each box of distribution represents the test set performance of 9 re-trains (three-fold, three random initialization each fold). The CNN model adopted model B as in Supplementary Fig 8. Abbreviations: AMC (Amoxicillin-Clavulanate), AMK (Amikacin), AMP (Ampicillin), AMX (Amoxicillin), CET (Cephalothin), CIP (Ciprofloxacin), CLI (Clindamycin), CPM (Capreomycin), CSN (Cycloserine), CTX (Cefotaxime), CTZ (Ceftazidime), CXM (Cefuroxime), EMB (Ethambutol), ERY (Erythromycin), ETH (Ethionamide), FUS (Fusidic Acid), GEN (Gentamicin), INH (Isoniazid), KAN (Kanamycin), MET (Methicillin), MXF (Moxifloxacin), OFL (Ofloxacin), PAS (Para-Aminosalicylic Acid), PEN (Penicillin), PZA (Pyrazinamide), RIF (Rifampicin), STR (Streptomycin), TBM (Tobramycin), TET (Tetracycline), TMP (Trimethoprim). The sample size is listed in brackets next to each phenotype predicted, format as amount of the “(resistant/susceptible)”. Dashed horizontal lines stand for AUROC of 0.8 (lightgrey) and 0.9 (pink).

## Discussion

In this study, we presented a novel manner in studying bacterial pan-genome by evaluating g2g predictability. The high average predictability among large amount of strains and genes demonstrated the generalizability. This g2g correlation allowed strain characteristics representation using very limited genomic information, which was tested with in vitro resistance prediction (Figure 5). While gene co-occurrence^10–12^ and LD^23–26^ only considered binary states of genes, g2g correlation further treated each gene as a multi-state object, which allowed correlation analysis among complex patterns (Supplementary Fig 18).

The multi-state correlation exhibited high complexity and the predictability lasted over a long distance. Taking the gene *gltD*, which ranked 12^th^ in CVL of *E. coli,* as an example, we predicted its SVs with only the 2^nd^-ranked gene *mrcB* or only the 3^rd^-ranked gene *metE*, which were 1.50±0.22 Mbp and 0.66±0.10 Mbp away from *gltD*, respectively (Supplementary Data 14, Supplementary Fig 19a). Although both input genes showed high prediction accuracy for most SVs of the *gltD*, the *mcrB* sometimes miss-predicted between SV3 and SV6, and the *metE* completely miss-predicted between SV2 and SV6. Miss-predictions do not result from high sequence similarity among SV2, 3 and 6 of *gltD*, since SV4 was also highly similar with SV2 and SV6 but was rarely miss-predicted (Supplementary Fig 16, 19a). In Supplementary Fig 19b, those strains with *gltD* of SV2 and 6 shared one same SV of *metE*, which explained those miss-predictions when *metE* was the single input. The observed correspondence mode was more than simply one-to-one among SVs, which was rarely explored in biallelic association studies of LD.

The correlation in our study could maintain stable over millions of bps, we believe this long distance predictability resulted from clonal structure embedded in our datasets. High g2g predictability could reach the farthest position in a circular chromosome for any two genes (Supplementary Fig 19c)^43–45^. Differently, LD decay in previous studies^23–26^ showed steep initial reduction and maintained a genome-wide low non-zero linkage. This difference resulted from our dataset design, which consists of clonal strains within each cgST. On the contrary, LD calculation usually selects very limited amount of representatives from each linage to account for the clonal structure of bacteria. We believe g2g correlation was formed in a process that genes co-evolved and experienced co-selection as well as clonal inheritance under environmental pressure. To adapt to a given environment, the whole genome could synchronously evolve because pressure on bacteria came from multiple aspects. Genes were screened to be present, absent and formed different SVs to better adapt to multi-faceted pressure, therefore combinations of gene status were gradually fixed by dominant clones. This process did not required direct functional interaction among genes, but generations passage in the same environment finally led to strong g2g correlation and predictability.

Notably, while many genes had expectedly high g2g predictability in such a clonal dataset, many others with low predictability were suggested to be very unclonal during their inheritance, regardless of core or accessory (Supplementary Fig 20a). We investigated factors associated with high or low predictability. However, neither gene presence frequency nor CVL values of genes did not showed a significantly correlation with g2g predictability (Supplementary Fig 20a, b, Supplementary Data 14). Categorizing gene tropism without bias remains challenging due to the lack of reliable pathway or functional annotations for many pan-genes. Further studies are required to elucidate the variability in predictability.

The strong correlation among genes demonstrated that genotyping has a large pool of feasible genes and potential in phenotype inference. For example, *E. coli* cgST B2 could be identified by absence of *espX4* and *espL1* without whole genome sequencing and complicated bioinformatics processes. New typing strategies can be independent of reference strains requiring a small CNN or RF model trained on a moderate scale of sample size and providing better genome representation. Our study proposed a set of algorithms and methods to test and find limited core genes, which allowed the representation of a genome. By embedding with pre-trained model, the representation of a long DNA sequence from genome can be further reduced to a vector of less than a thousand float numbers. All these efforts will bring feasibility and efficiency for the future large scale analysis.

It is worth noting that k-mer patterns from either high or low CVL input combinations could predict cgST with high accuracy (Supplementary Fig 21, 22), but within each cgST, k-mer patterns from different input combinations exhibited significant differences (Supplementary Fig 23, Supplementary Data 15). This means patterns learned by models were not generalizable among different input windows, and this was also the reason we evaluated input windows by re-train based cross validations. With models pre-trained on complete core genome, we found that performance of re-prediction using low CVL input window failed to maintain a high accuracy similar as re-train results (Supplementary Fig 24 first row, Figure 3). Pre-trained models even with powerful structures like Res-Net^46^ could not properly reflect the accuracy of low CVL input windows in inferring the status of other genes (Supplementary Fig 25 and Supplementary Fig 24 second row).

Last but not least, it is important to acknowledge that the genes analyzed in this study represent a small subset of the entire pan-genome. In *E. coli* Dataset II, a total of 72,275 gene clusters were identified, similar as the previous study^4^. Notably, 61,051 of these clusters were found in only one strain each, indicating that a lot of genes lack enough presence and prevalent SVs in *E. coli* SAC. Therefore, the status predictability of many pan-genes remained unknown and require much larger dataset in the future to explore their g2g correlation.

## Methods

### *E. coli* isolation from UTI patients and sequencing

From First Affiliated Hospital of Kunming Medical College, middle stage urine samples from within 2 hours post micturition were collected and 100 μL were plated on 90 mm MacConkey Agar (Antu Biotechnology, Zhengzhou), 37°C for 8 hours. 1000 single colonies and more led to diagnosis of UTI patients whose samples were included. Plates were transferred under −4°C to laboratory of Center of Life Science, Yunnan University, and samples were assigned a unique identifier and subjected to re-cultivation using 20mL LB culture (Solarbio, Beijing, PM0011-500g), 37°C over night under 220 rpm. To ensure long-term viability, 700 μL resultant bacterial cultures were combined with an equal volume of glycerol (Sangon Biotech, Shanghai, A600232-0500) and stored under appropriate conditions.

A 10μL aliquot of the culture was aseptically sampled using a sterile loop and inoculated into 5mL LB/MHB (Solarbio, Beijing, M8556) culture, followed by incubation with agitation at 220 rpm, 37°C over night. Subsequently, 100μL of this enriched culture was transferred to 5 mL fresh LB/MHB for subculture, shaked under 225 rpm, 37°C or 4 h. Serial dilution was then performed on the subcultured samples, achieving a 10^−3^ dilution factor, and the diluted cultures were plated onto 90 mm LB plates for 37°C over night.

*E. coli* Colonies were picked according to morphology and selected colonies were further propagated in 5mL LB/MHB to enrich cell populations for downstream analysis. Genomic DNA was extracted from these enriched cultures using a commercial DNA extraction kit (Beyotime Biotechnology, Shanghai, B518225), adhering strictly to the manufacturer’s protocols to ensure high-quality DNA for sequencing purposes.

Species identification was performed using full length 16S ribosomal RNA gene sequencing. Libraries were constructed for identified *E. coli* strains to perform paired-end 150 bp sequencing using NovaSeq 6000 (Major Biotechnology, Shanghai).

### Minimum Inhibitory Concentration (MIC) determination

MIC values are defined as the lowest concentration that results in a more than 90% reduction in bacterial growth, or the absence of well-defined bacterial spheres. 50μL MHB was added to the 96-well plate to dilute antibiotics, then 50 L (1×10^5^ CFU/mL) bacterial suspension was added to each well, mixed, and incubated at 37°C for 18h. Bacterial growth is determined by measuring the optical density at a wavelength of 600nm (OD600) (Thermofisher, Shanghai, A51119600). MIC test of this study included Kanamycin (BBI, Shanghai, A600286), Fosfomycin (Northeast Pharmaceutical Group, Liaoning, 4210509), Cephalothin (Solarbio, Beijing, C8500), Nitrofurantoin (Tianjin Lisheng Pharmaceutical, Tianjin, 2106015).

### PCR target gene detection

Frozen stored strains YN-24, YN-9, YN-91, YN-16, YN-54, YN-66, YN-57, YN-35, YN-48, and YN-89, were streaked with LB. Incubate strains on plates at 37°C for 12 hours to promote growth. Following incubation, isolate a single colony from each plate and inoculate it into 5 mL of LB broth. Then, incubate these cultures at 37°C with a shaking speed of 220 rpm for 10 hours to ensure optimal bacterial growth.

For genomic DNA extraction, utilize TIANGEN’s Bacterial DNA Kit (DP302-02, Beijing) to purify the total bacterial DNA from the cultivated strains. Adjust the concentration of the purified DNA to 10 ng/μL for subsequent polymerase chain reaction (PCR) analyses. Prepare the PCR reaction mix as follows: 5μL of 2x Rapid Taq Master Mix (Vazyme, Nanjing P222-03), 3.2 L of ddH O, 1 L of template DNA, and 0.8μL of the designated primers. The PCR amplification protocol consists of an initial denaturation at 95°C for 3 minutes, followed by 30 cycles of denaturation at 95°C for 15 seconds, annealing at 60°C for 15 seconds, and extension at 72°C for 30 seconds, concluding with a final extension step at 72°C for 5 minutes. The specific primers used in this study are as follows: for *espX4*, forward primer CATGGTGGGTGAAGGA and reverse primer CGATTAGCTCGCCACTC; for *espL1*, forward primer CGATGCTGTCATACTGTG and reverse primer GAGAGGAACCGTGACA.

### Bioinformatics

Sequencing results of lab isolated strains and downloaded SRA (Sequence Read Archive) reads, were quality controlled using fastp (v0.23.2)^47^, where parameters were adjusted according to read length and adapter length. Including read pairs with enough quality, reads with only one direction passed the filter were also included in the assembly construction using SPAdes (v3.15.5)^48^.

Bulk assemblies data downloading for SAC construction was performed using customized workflow blast_at_local_computer^49^, which transformed GCA accession into rsync address using Biopython (v1.83)^50^ and downloaded data using rsync from NCBI assembly database. The process included automated md5sum check and re-download of failed ones. The downloaded assemblies were primarily quality checked using QUAST (v5.2.0)^51^ to filter those with N50 < 200,000 bp. The phenotype dataset were selected from BV-BRC^41^ and assemblies were downloaded. Samples whose assembly not available from BV-BRC were obtained from SRA using sratoolkit(v3.0.5)^52^.

Downloaded assemblies were randomly selected for construction of Dataset I and II. These selected assemblies were quality checked using checkM (v1.2.2)^53^ Taxonomic-specific Workflow, in which marker file were generated for each species and then the completeness and contamination rate were assessed. Assemblies with completeness <95% and contamination rate >5% were excluded from construction of Dataset I and II.

All assemblies were re-annotated using Prokka (v1.14.6)^54^ under fast mode which only detect genes without function annotation. The annotated DNA sequences including coding region, tRNA and non-coding RNA sequences were further clustered by customized pipeline OrthoSLC (v0.2.4). It utilize 100% similarity threshold pre-clustering and (Python or C++ built in) hash algorithm for binning to highly accelerate the all-vs-all blast, RBBH pair screening, reciprocal best finding and single linkage clustering. This allowed the gene cluster construction of 801 *L. monocytogenes* and 1200 *E. coli* assemblies could be finished within 25 and 320 minutes using 36 threads, respectively. The RBBH pair was screened using a limit of length difference >0.9, which means the shorter sequence in the pair shall not be shorter than 90% of the longer one. The clusters with exact 1 gene from each assembly were defined as StCGCs.

Multi-sequence alignment on the StCGCs were performed using Kalign (v3.3.5)^55^ default mode and aligned clusters were later concatenated into core genome of each strain using customized script. The phylogeny computation utilized RAxML (v8.0.2)^56^ under GTRCAT mode. 1000 rapid bootstraps were performed on the phylogenetic tree in Figure 1 while all other trees included in the study performed only 200 rapid bootstraps. The cgST identification was assisted by reference assemblies and followed ‘one node one cgST’ rule. All strains classified as a certain cgST should share a common node in the phylogeny structure and cgSTs could be identified from a given cgST using sub-nodes.

The VFs annotation on Dataset I, II and SAC were achieved by blastn (v2.13+) VFDB (Feb 10 2023) DNA sequences core dataset (setA)^32^ against assemblies. Every best hit in the results were followed by screening that the aligned region should reach >80% similarity with reference and length difference limit between hit region and reference should be >0.8. For 91 patient isolated *E. coli*, the hit regions on assemblies were further tested coverage using bowtie2 (v2.5.0)^57^. The ARGs annotation on 137 *E. coli* in Figure 1 and Dataset II were achieved by inputing assemblies into Card (3.0.2) and Resistance Gene Identifier (RGI) (v6.0.2)^33^. And gene resistance mechanism for each gene was obtained by RGI heatmap generation function. ARGs annotation on SAC utilized the ARGs found in Dataset II of each species, and blastn (v2.13+)^58^ these genes against each assembly, with same >80% similarity and >0.8 length difference degree filtering.

Function annotation on each sequence of StCGCs was performed using eggNOG-mapper (v2.1.12) and eggdb (v5.0.2)^35,36^, in which DIAMOND (v2.0.11.149) was employed for searching. The function annotation of each cluster was summarized and heterofunctional cluster was defined as cluster containing at least 2 kinds of function, where the function with least sequences in the cluster shall have more than _limit_ sequences let a cluster become heterofunctional. Where:

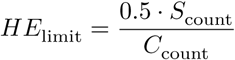

*S_count_* and *C_count_* stand for the count of strains and count of cgSTs, in Dataset II of given species, respectively. This set up is dependent on the even distribution of samples in cgSTs, so that if a StCGC contained a different function with amount of sequences more than half of one cgST strain amount, the determination of one of the cgSTs had possibility to be decided by this hterofunctional cluster. In other words, if a StCGC of *E. coli* (*S*_count_ = 1200, *C*_count_ = 6, *HE*_limit_ = 100) contained only 10 genes that were of different function from other 1190 sequences, this cluster could not differ a cgST from other simply by its function composition.

The Cluster Variation Level (CVL) was calculated on each StCGC:

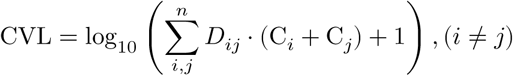

where *n* stands for all pairwise combinations of dereplicated unique SVs in a give cluster; *D_ij_* stands for the count of different sites in the alignment of unique sequence *i* and *j*; *C_i_* and *C_i_* stand for the amount of unique sequence *i* and *j* in the cluster. The algorithm attempts to value the variation level within a cluster and clusters were then ranked according to their CVL value for input window design.

To extract core genes of StCGC (Figure 4) and pan-genes (presence frequency >5% and <100% in Dataset II of *E. coli*, Supplementary Fig 14 second column) from SAC, the most frequent SV in each of gene cluster of Dataset II were deemed as seeds. After blastn (v2.13+)^58^ these seeds against SAC assemblies, each best hit in an assembly with >0.9 and >0.8 length difference degree were accepted for core genes and pan-genes, respectively.

Gene extracted from SAC were grouped using Python3 built in hash algorithm for dereplication and SV summary. All sequences extracted was checked using customized scripts to ensure no hash collision occurred. The SV summary was later used for sequence presence, SV prediction labels. Distance among StCGCs were calculated using complete assemblies in Supplementary Table 1.

k-mer frequency was extracted using jellyfish (v2.3.0)^59^. The corrected frequency of a single k-mer was obtained by rescaling the frequency to a range between 0 and 1, by reduction of the lowest frequency of this k-mer among all samples and division by the highest frequency after reduction.

Task parallelrization was achieved through customized Python3 scripts and GNU parallel^60^ for efficiency.

### Machine learning

LR based cross validation, which was applied in cgST prediction using k-mer, VFs pattern, and ARGs pattern as input, was performed in R (v4.3.1)^61^ with glmnet (v4.1-8)^62^ package. Each re-train started with randomly selected training set (70% of each cgST). An extra re-train would be added if a re-train did not reach convergence until wanted count of re-train was met. Final average AUROC and average weight of each input feature were calculated to evaluate the performance and to pick features significant for the prediction.

The RF model used in pseudo-labeling based dataset construction was R randomforest (v4.7-1.1) package^63^, where the number of trees used was 2 fold of the input dimension. The trained models for pseudo-label prediction was tested to have >98% test set (30% samples from each cgST of Dataset I) performance before actual prediction.

The CNN models were constructed using Pytorch (v2.0.0)^64^ and DNA sequences were transformed into usable matrix with a one-hot like encoding manner as in Supplementary Fig 7. Input DNA sequences were first transformed using a one-hot like manner. The transformation put IPUAC DNA ambiguity into considerations. Model A were for cgST prediction which included 4 convolution layers and 4 maxpooling layers and only the first maxpooling and convolution layers had 6 channels. Model A had a deeper structure than model B due to different input size. Model A took complete core genome size as input (∼346,000 bp to 890,000 bp), therefore, we designed a deeper structure for feature extraction and dimension reduction. While model B took partial genome (∼14,000 bp to 27,000 bp) as input to predict accessory gene presence and SV of other genes. Model B had only 2 convolution layers and 2 maxpooling layers and the amount of channels for first convolution and maxpooling layers was equal to the count of class (SVs) to be predicted. The channel count in model B was set to 6 in phenotype prediction.

The evaluation of cgST and phenotype prediction using different input combinations was through 9 re-trains, containing 3 fold cross validation and 3 random initialization each fold. Each fold contained nearly equal amount of samples and strains of each class were evenly distributed in 3 folds. The evaluation of accessory gene presence and SV prediction was evaluated using 4 re-trains, 2 fold and 2 random initialization each fold. In sequence prediction, the gene to be predicted had at least 5,000 presence and absence in SAC, each fold contained 2,500 samples with target genes present in their assemblies and another 2,500 samples without. In SV prediction, assemblies owning the exactly identical sequences will be grouped as owning the same SV, equivalent to clustering of 100% similarity. Genes whose SV was predicted had at least 2 prevalent SVs that had at least 2,000 presence in SAC. The prediction would include a class ‘other’ that containing all non-prevalent SVs, and if this class ‘other’ also had >2,000 presence in SAC. Each of the 2 folds of cross validation contained 1,000 samples of each SVs. The performance evaluation of binary employed AUROC and multiple classification employed accuracy, which was calculated as the correct predictions out of all predictions since even class count was ensured.

The CNN models employed the learning rate of 3×10^−3^ and 5×10^−3^ with 150 epoches in prediction using model A and B, respectively. The batch size design for training was 200 in cgST prediction, 2,000 in accessory gene presence prediction, and one fourth of training set size in SV prediction (to adapt 1,000 more samples every 1 additional class). For phenotype prediction, the learning rate was 1×10^−3^ with batch size of 120 for 600 epoches. To select the best model among training epoches, the epoche with lowest test set loss was chosen for prediction of cgST, accessory gene presence, and SV prediction; and in phenotype prediction, the epoche with highest AUROC was selected. Notably, train set performance of each epoch was represented by the first batch before gradient descent. Loss function of Cross Entropy Loss (CrossEntropyLoss) and Binary Cross Entropy Loss (BCELoss) were employed for binary and multi-class classifications, respectively.

The embedding was performed using two separately trained model for input of 1 ∼ 6 (CVL rank) and 201 ∼ 218, respectively. Structure of model B was trained to predict StCGC with ID “10207-13847” in *E. coli* (Supplementary Data 7), which had 10 prevalent SVs. The pre-trained model was the on with lowest loss over 150 training epoches, and both input of 1 ∼ 6 and 201 ∼ 218 were padded to the same length to use same model hyper-parameters. Using the pre-trained models, all input were pre-calculated until the final layer to achieve DNA to vector (DNA2vec) embedding. In pan-genes status predictions, these vectors became the input and only the final full connection layer was trained to predict each gene.

For re-prediction based evaluation of cgST prediction accuracy using core gene combinations, pre-trained model A and res-net like model were generated using 70% randomly selected samples from each cgST. Res-net like model starts with 2 convolutional layers for a primary dimension reduction and included ReLu and batch normalization which was not included in model A. Both models were trained for 500 epoches and learning rate of 3×10^−3^. While model A took batch size of 210, res-net like model took batch size of 80 and a L2 regularization of 0.1 to entourage the learning of more features. 10 Models of model A and res-net, respectively, with >98% test set performance were kept for re-prediction. The re-prediction input included both training and test set samples used in pre-trained model generation.

### Statistics

Principle Coordinate Analysis (PCoA) was performed using R vegan (2.6-4)^65^ package for VFs pattern, ARGs pattern and k-mer pattern within cgST analysis. Significance test pairwise among groups were performed using PERMANOVA algorithm with 999 permutations. Data visualization in this study was facilitated by R ggplot2 (v3.4.3)^66^, ggtree (v3.10.0)^67^ packages.

### Software, workflow management

The pipeline and softwares were managed using Docker (v>4.16).

## Code availability

Bulk assembly downloading pipeline is available at https://github.com/xchuam/blast_at_local_computer; Gene clustering pipeline OrthoSLC (v0.2.4) is available at https://github.com/JJChenCharly/OrthoSLC; All machine learning, bioinformatics and partial visualization codes are available at https://github.com/JJChenCharly/g2g.

## Data availability

The Data will be made available upon request.

## Declaration of Competing Interest

The authors declare no competing interests.

## Ethnicity statement

All procedures involving the collection and analysis of strains from UTI patients were carried out in compliance with the ethical standards of the institutional and/or national research committee. The study protocols were approved by the Ethics Committee of The First Affiliated Hospital of Kunming Medical College (2022L203) and The Committee on Human Subject Research and Ethics, Yunnan University (CHSRE number: CHSRE2022030). Informed consent was obtained from all individual participants included in the study.

## Acknowledgments

This work was supported by the National Natural Science Foundation of China (32070818 and 82460142 to C.M.G.), Natural Science Foundation of Yunnan Province of China [202001BB050005], the Xingdian talent support program of Yunnan Province. Guangdong Hybribio Biotech Funding (H20230313). We thank Songnian Hu (University of Chinese Academy of Science) for early stage conceptualization support; Cheng Peng (Yunnan University) for bioinformatics and computation hardware support; Longyu Chen and Zirui Wang (University of Chinese Academy of Science) for machine learning consultation and computation power support.

## Supplementary Tables

**Supplementary Table 1:**
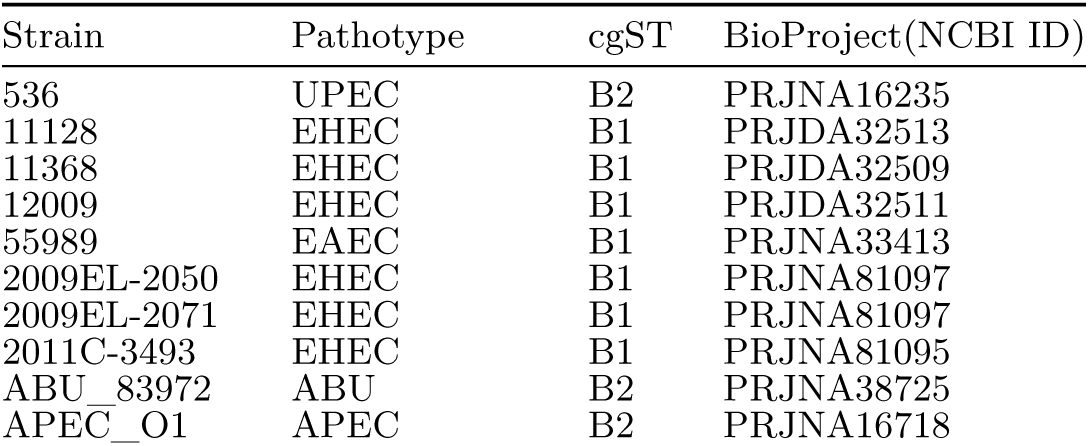
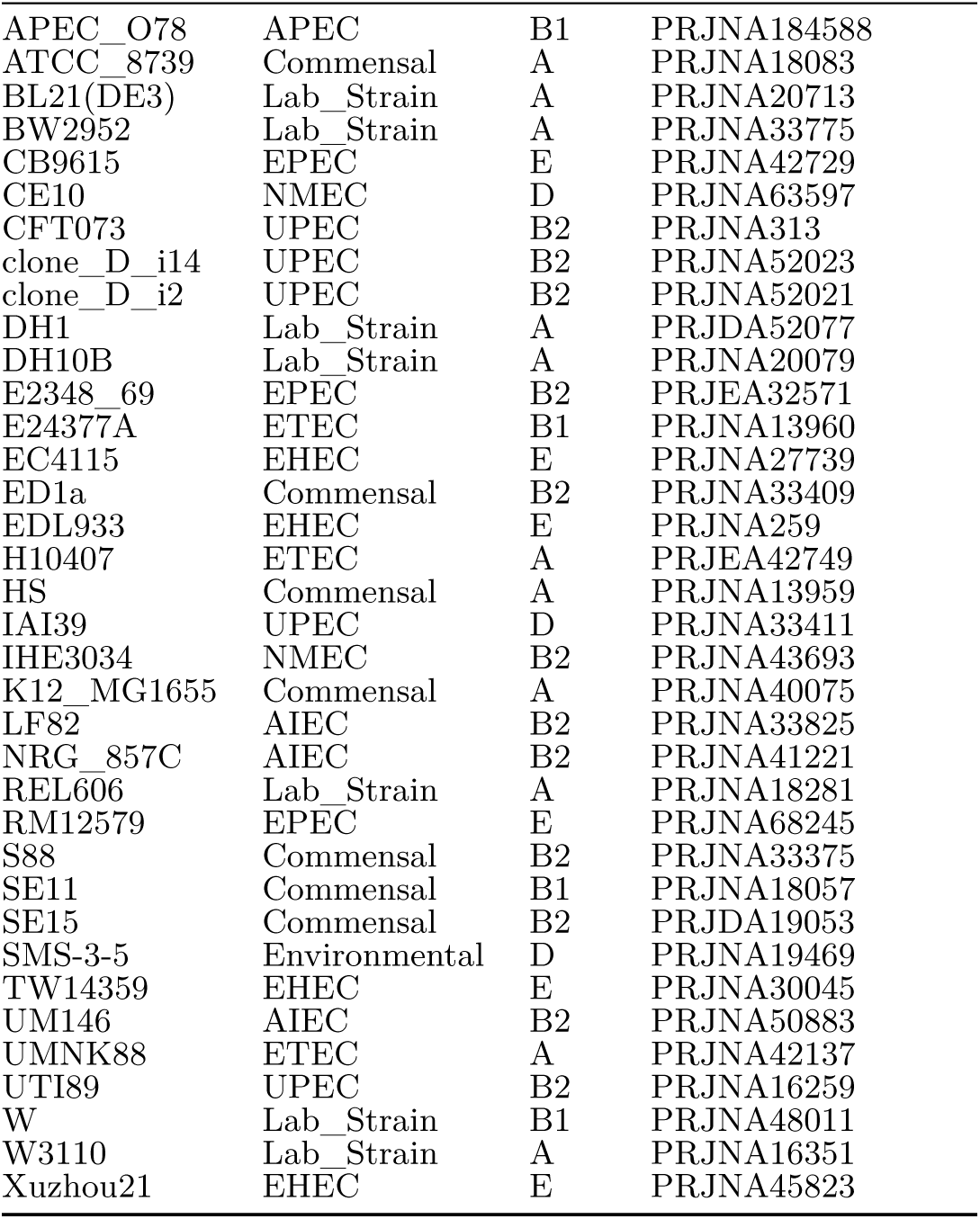
46 reference *E. coli* genome information. The genome included the circular chromosome and plasmids. These reference strains were used for cgST identification in Fig 1 and 2 for *E. coli* dataset.

**Supplementary Table 2:**
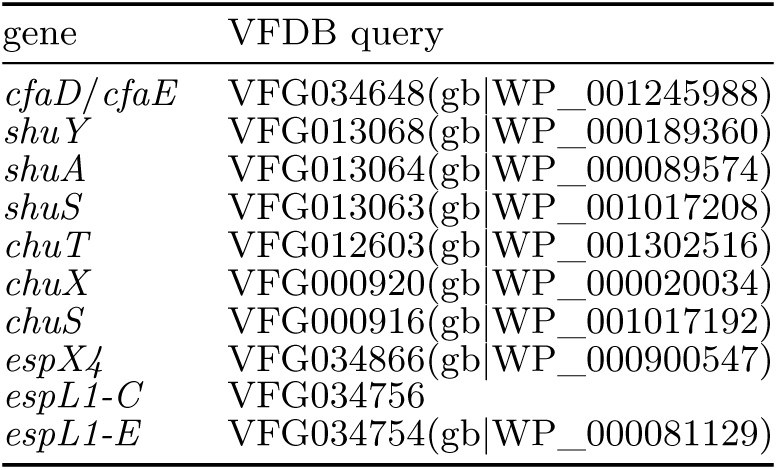
Gene name and VFDB ^1^ query correspondence. *espL1-C* and *espL1-E* stands for *espL1* gene from *E. coli* CB9615 and EDL933, respectively. The query accession could be used to search relavent information (related literature, ortholog, mechanisms) from VFDB.

| gene | VFDB query |
| --- | --- |
| *cfaD/cfaE* | VFG034648(gb\|WP_001245988) |
| *shuY* | VFG013068(gb\|WP_000189360) |
| *shuA* | VFG013064(gb\|WP_000089574) |
| *shuS* | VFG013063(gb\|WP_001017208) |
| *chuT* | VFG012603(gb\|WP_001302516) |
| *chuX* | VFG000920(gb\|WP_000020034) |
| *chuS* | VFG000916(gb\|WP_001017192) |
| *espX4* | VFG034866(gb\|WP_000900547) |
| *espL1-C* | VFG034756 |
| *espL1-E* | VFG034754(gb\|WP_000081129) |

**Supplementary Table 3:** cgST reference strains. Strains selected in Dataset I to be cgST identification reference in phylogeny in Dataset II of *E. coli*.

| ID in Dataset II | GCA | Description |
| --- | --- | --- |
| EC_F_ref0 | GCA_014794105.1 | GCA_014794105.1_PDT000848429.1_genomic |
| EC_F_ref1 | GCA_000459215.1 | GCA_000459215.1_Esch_coli_HVH_208_4-3112292_V1_genomic |
| EC_F_ref2 | GCA_021425225.1 | GCA_021425225.1_PDT001218343.1_genomic |
| EC_F_ref3 | GCA_021085865.1 | GCA_021085865.1_PDT001197641.1_genomic |
| EC_F_ref4 | GCA_019204805.1 | GCA_019204805.1_PDT001088470.1_genomic |

**Supplementary Table 4:** cgST reference strains. Strains selected in Dataset I to be cgST identification reference in phylogeny in Dataset II of *L. monocytogenes*.

| ID in Dataset II | GCA or GI | Description |
| --- | --- | --- |
| LM_A_ref0 | GCA_020988225.1 | GCA_020988225.1_PDT001185688.1_genomic |
| LM_A_ref1 | GCA_003685255.1 | GCA_003685255.1_PDT000365297.1_genomic |
| LM_A_ref2 | GCA_004598375.1 | GCA_004598375.1_PDT000218577.2_genomic |
| LM_A_ref3 | GCA_009449765.1 | GCA_009449765.1_PDT000421349.2_genomic |
| LM_A_ref4 | GCA_024119015.1 | GCA_024119015.1_PDT001338036.3_genomic |
| LM_B_ref0 | GCA_004484765.1 | GCA_004484765.1_PDT000065844.2_genomic |
| LM_B_ref1 | GCA_018225825.1 | GCA_018225825.1_PDT001017255.1_genomic |
| LM_B_ref2 | GCA_028162645.1 | GCA_028162645.1_PDT001589962.1_genomic |
| LM_B_ref3 | GCA_023538895.1 | GCA_023538895.1_PDT001314493.1_genomic |
| LM_B_ref4 | GCA_009490045.1 | GCA_009490045.1_PDT000454753.2_genomic |
| LM_C_ref0 | GCA_020757795.1 | GCA_020757795.1_PDT001168515.1_genomic |
| LM_C_ref1 | GCA_000307065.1 | GCA_000307065.1_ASM30706v1_genomic |
| LM_C_ref2 | GCA_010080955.1 | GCA_010080955.1_PDT000626992.1_genomic |
| LM_C_ref3 | GCA_004472405.1 | GCA_004472405.1_PDT000447650.1_genomic |
| LM_C_ref4 | GCA_009361515.1 | GCA_009361515.1_ASM936151v1_genomic |
| 10403S | GI: 345535581 | cgST D |
| EGD | GI: 543859294 | cgST D |
| EGD-e | GI: 1796433495 | cgST D |
| LO28 | GI: 2173384332 | cgST D |
| Scott_A | GI: 1796436383 | cgST B |

**Supplementary Table 5:** cgST reference strains. Strains selected in Dataset I to be cgST identification reference in phylogeny in Dataset II of *C. jejuni*.

| ID in Dataset II | GCA | Description |
| --- | --- | --- |
| CJ_B_ref0 | GCA_018994985.1 | GCA_018994985.1_PDT001074550.1_genomic |
| CJ_B_ref1 | GCA_008248845.1 | GCA_008248845.1_PDT000573498.1_genomic |
| CJ_B_ref2 | GCA_005373965.1 | GCA_005373965.1_PDT000370886.1_genomic |
| CJ_B_ref3 | GCA_017182455.1 | GCA_017182455.1_PDT000945415.1_genomic |
| CJ_B_ref4 | GCA_010299745.1 | GCA_010299745.1_PDT000563316.1_genomic |
| CJ_C_ref0 | GCA_004822525.1 | GCA_004822525.1_PDT000406612.1_genomic |
| CJ_C_ref1 | GCA_004987965.1 | GCA_004987965.1_PDT000271318.2_genomic |
| CJ_C_ref2 | GCA_028591385.1 | GCA_028591385.1_PDT001612276.1_genomic |
| CJ_C_ref3 | GCA_005316785.1 | GCA_005316785.1_PDT000266958.5_genomic |
| CJ_C_ref4 | GCA_009188145.1 | GCA_009188145.1_PDT000610439.1_genomic |
| CJ_D_ref0 | GCA_004278335.1 | GCA_004278335.1_PDT000384529.1_genomic |
| CJ_D_ref1 | GCA_008170205.1 | GCA_008170205.1_PDT000570024.1_genomic |
| CJ_D_ref2 | GCA_025939745.1 | GCA_025939745.1_ASM2593974v1_genomic |
| CJ_D_ref3 | GCA_017239835.1 | GCA_017239835.1_PDT000939616.1_genomic |
| CJ_D_ref4 | GCA_011737495.1 | GCA_011737495.1_PDT000712334.1_genomic |
| CJ_A_ref0 | GCA_005354025.1 | GCA_005354025.1_PDT000360883.1_genomic |
| CJ_A_ref1 | GCA_025020425.1 | GCA_025020425.1_PDT001409990.1_genomic |
| CJ_A_ref2 | GCA_005156375.1 | GCA_005156375.1_PDT000173386.2_genomic |
| CJ_A_ref3 | GCA_015088835.1 | GCA_015088835.1_PDT000856263.1_genomic |
| CJ_A_ref4 | GCA_004903555.1 | GCA_004903555.1_PDT000428963.1_genomic |

**Supplementary Table 6:**
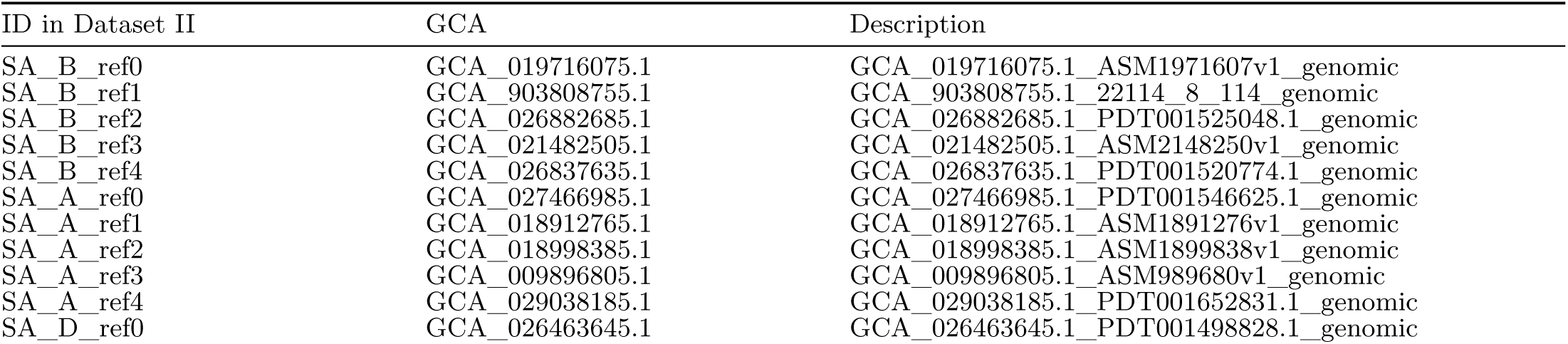
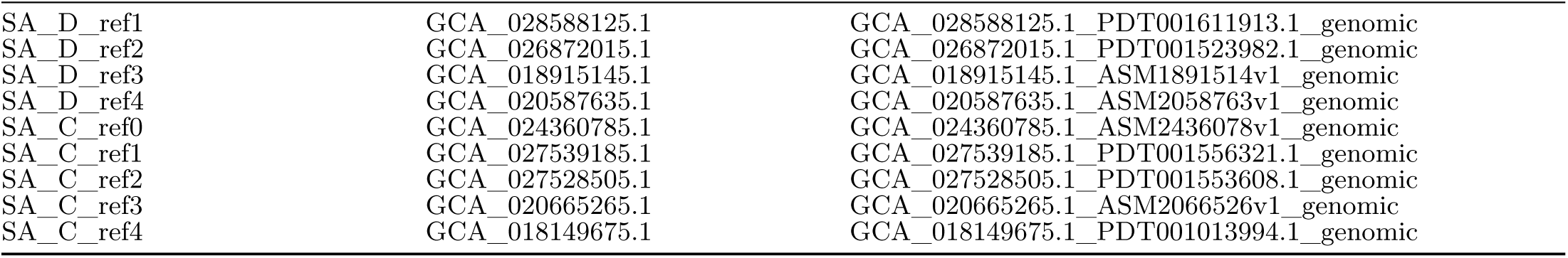
cgST reference strains. Strains selected in Dataset I to be cgST identification reference in phylogeny in Dataset II of *S. aureus*.

**Supplementary Table 7:** Studies and species included from the study. Studies were selected from BV-BRC^2,3^ from which assemblies were obtained. Corresponding strain ID and resistance label to each antibiotics are in Supplementary Data 11.

| Study | Species included |
| --- | --- |
| Davies et al.<sup>4</sup> | *E. coli* |
| Moradigaravand et al.<sup>5</sup> | *E. coli* |
| Bradley et al.<sup>6</sup> | *S. aureus* and *M. tuberculosis* |
| Kumar et al.<sup>7</sup> | *S. aureus* |
| Coll et al.<sup>8</sup> | *M. tuberculosis* |

**Supplementary Table 8:** House keeping genes. The house keeping genes selected from published datasets. Their names, total length of target regions for E. coli^9^, L. monocytogenes^10^, C. jejuni^11^, S. aureus^12^, and M. tuberculosis^13–15^

| Species | house keeping genes | total length |
| --- | --- | --- |
| *E. coli* | *adk, fumC, gyrB, icd, mdh, purA, recA* | 3423 bp |
| *L. monocytogenes* | *abcZ, bglA, cat, dapE, dat, idh, ihkA* | 3288 bp |
| *C. jejuni* | *aspA, glnA, gltA, glyA, pgm, tkt, uncA* | 3309 bp |
| *S. aureus* | *arcC, aroE, glpF, gmk, pta, tpi* | 2670 bp |
| *M. tuberculosis* | *gyrA, gyrB, katG, purA, recA, rpoB, sodA* | 4612 bp |

## Supplementary Data

**Supplementary Data 1:** VFs, ARGs annotation and cgST of 46 reference and 91 patient isolated *E. coli*.

| cgST | VFs annotation | ARGs annotation |
| --- | --- | --- |
| cgST_137_Ec.csv | VFs_137_Ec.csv | ARGs_137_Ec.csv |

**Supplementary Data 2:**
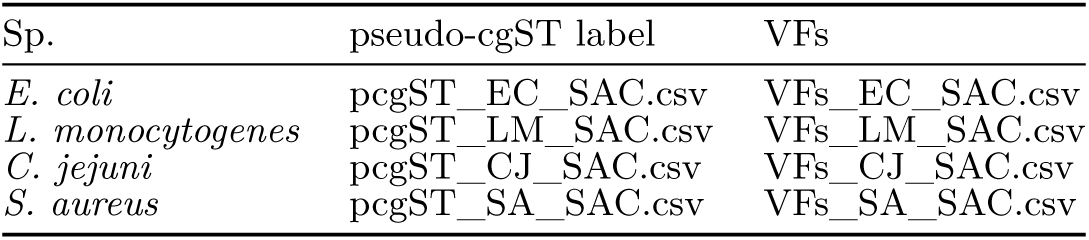
VFs and pseudo-cgST-label of SAC (Species Assembly Collection) of 4 species.

| Sp. | pseudo-cgST label | VFs |
| --- | --- | --- |
| *E. coli* | pcgST_EC_SAC.csv | VFs_EC_SAC.csv |
| *L. monocytogenes* | pcgST_LM_SAC.csv | VFs_LM_SAC.csv |
| *C. jejuni* | pcgST_CJ_SAC.csv | VFs_CJ_SAC.csv |
| *S. aureus* | pcgST_SA_SAC.csv | VFs_SA_SAC.csv |

**Supplementary Data 3:** VF annotation and cgST of Dataset I and II of 4 species.

| Sp. | cgST I | cgST II | VFs I | VFs II |
| --- | --- | --- | --- | --- |
| *E. coli* | cgST_EC1.csv | cgST_EC2.csv | VFs_EC1.csv | VFs_EC2.csv |
| *L. monocytogenes* | cgST_LM1.csv | cgST_LM2.csv | VFs_LM1.csv | VFs_LM2.csv |
| *C. jejuni* | cgST_CJ1.csv | cgST_CJ2.csv | VFs_CJ1.csv | VFs_CJ2.csv |
| *S. aureus* | cgST_SA1.csv | cgST_SA2.csv | VFs_SA1.csv | VFs_SA2.csv |

**Supplementary Data 4:** ARG annotation of Dataset II of 4 species.

| Sp. | ARGs II |
| --- | --- |
| *E. coli* | ARGs_EC2.csv |
| *L. monocytogenes* | ARGs_LM2.csv |
| *C. jejuni* | ARGs_CJ2.csv |
| *S. aureus* | ARGs_SA2.csv |

**Supplementary Data 5:** Average weight of 50 re-trains using VFs or ARGs to predict cgST in Dataset II.

| Sp. | ARGs weight | VFs weight |
| --- | --- | --- |
| *E. coli* | ARGs_wei_EC.csv | VFs_wei_EC.csv |
| *L. monocytogenes* | ARGs_wei_LM.csv | VFs_wei_LM.csv |
| *C. jejuni* | ARGs_wei_CJ.csv | VFs_wei_CJ.csv |
| *S. aureus* | ARGs_wei_SA.csv | VFs_wei_SA.csv |

**Supplementary Data 6:** VFs name (in Fig 3) and VFDB query correspondence; ARGs name (in Fig3) to CARD model name correspondence.

| ARGs | VFs |
| --- | --- |
| ARGs_CARD.XLSX | VFs_VFDB.XLSX |

**Supplementary Data 7:**
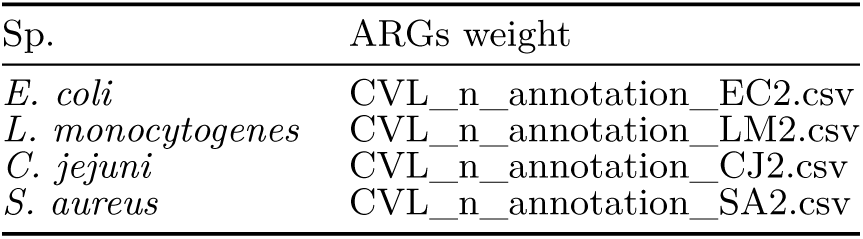
CVL result, StCGC cluster parameters, prokka/eggnogmapper annotation result, and the most prevalent sequence variant of StCGCs of Dataset II.

**Supplementary Data 8 (Fig 3): KO simulation for cgST recovery performance.** EC: *E. coli*, LM: *L. monocytogenes*, CJ: *C. jejuni*, SA: *S. aureus*.

**Supplementary Data 9:** Accessory gene presnece prediction results. EC: *E. coli*, LM: *L. monocytogenes*, CJ: *C. jejuni*, SA: *S. aureus*.

| Species | File |
| --- | --- |
| *E. coli* | presence_EC.xlsx |
| *L. monocytogenes* | presence_LM.xlsx |
| *C. jejuni* | presence_CJ.xlsx |
| *S. aureus* | presence_SA.xlsx |

**Supplementary Data 10:** Gene sequence variant prediction results. EC: *E. coli*, LM: *L. monocytogenes*, CJ: *C. jejuni*, SA: *S. aureus*.

| Species | File |
| --- | --- |
| *E. coli* | seqtype_EC.xlsx and seqtype_EC_other_excluded.xlsx |
| *L. monocytogenes* | seqtype_LM.xlsx |
| *C. jejuni* | seqtype_CJ.xlsx |
| *S. aureus* | seqtype_SA.xlsx |

**Supplementary Data 11: pan-gene status prediction results of *E. coli*.** presence and sequence variants prediciton result. The function annotation result includs the sequence of the most prevalent sequence variant and frequency in SAC.

**Supplementary Data 12: Phenotype labels for data from published datasets. Phenotype_labels.xlsx.**

**Supplementary Data 13: Phenotype prediction result. Phenotype_prediction_result.csv.**

**Supplementary Data 14:**
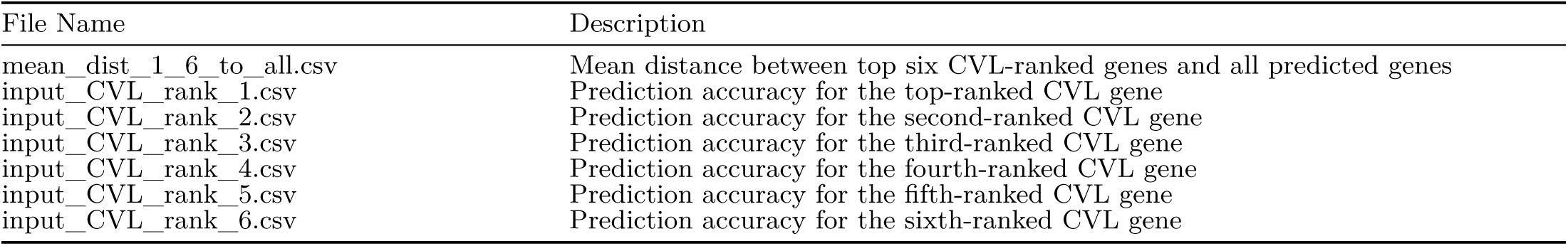
Distance table and prediction results by single gene of r Ec_‘ Dataset II. Prediction accuracy by each of top six CVL rank genes, and describingdistances between prediction input genes and genes predicted.

**Supplementary Data 15:** *E. coli* 3 to 7-mer frequency. file name format: “input gene CVL rank”_“k-mer length”-mer.csv.

| Input CVL rank | 3-mer | 7-mer |
| --- | --- | --- |
| 1-6 | 1-6_3-mer.csv | 1-6_7-mer.csv |
| 201-206 | 201-206_3-mer.csv | 201-206_7-mer.csv |
| 400-406 | 400-406_3-mer.csv | 400-406_7-mer.csv |
| odd | odd_3-mer.csv | odd_7-mer.csv |
| even | even_3-mer.csv | even_7-mer.csv |

## Supplementary Figures

**Supplementary Fig 1:**
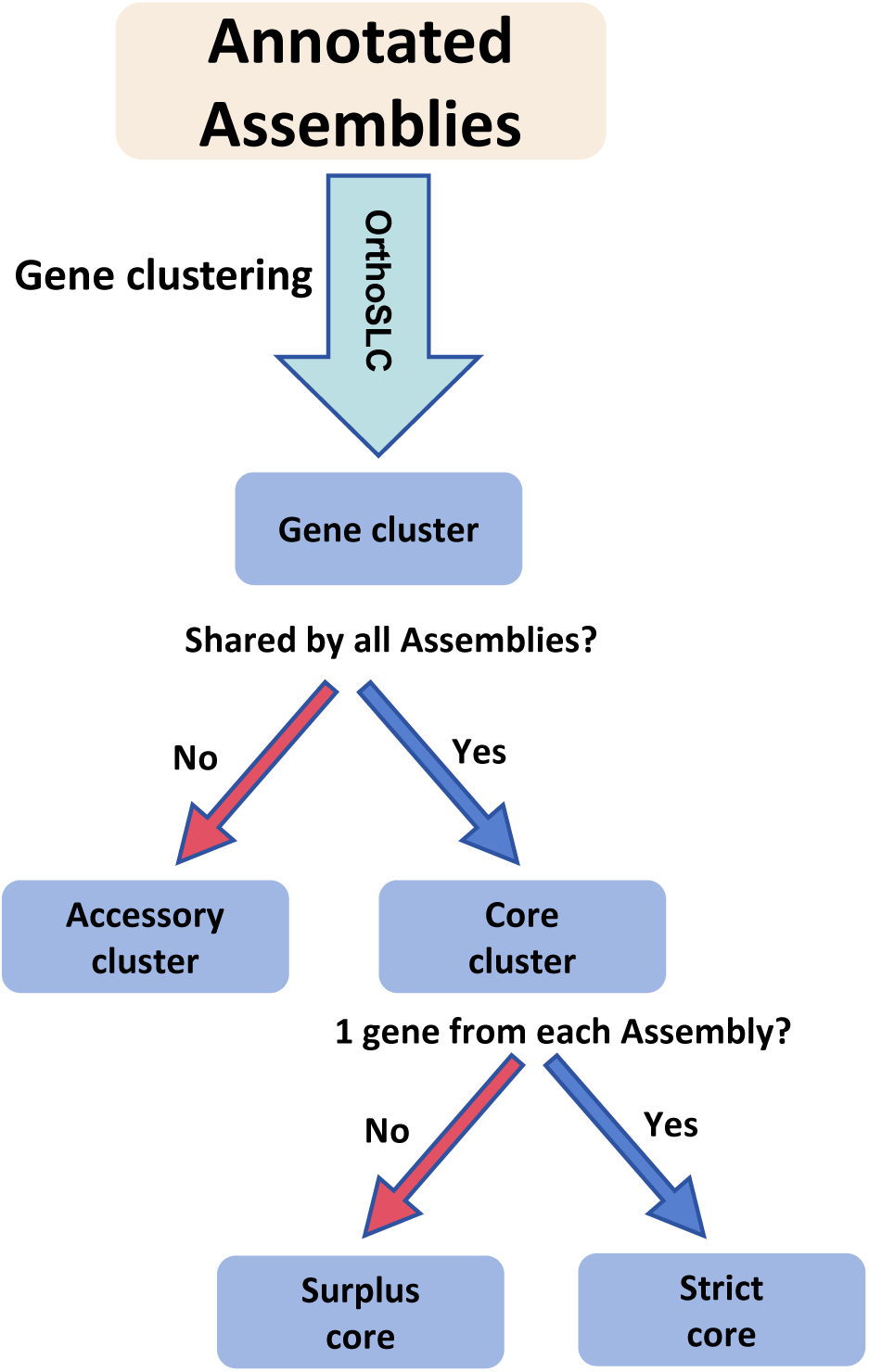
Categorization of gene clusters. Gene Clusters shared by only a subset of assemblies participated analysis are defined as accessory gene clusters. If a cluster had more number of genes than the number of assemblies participated analysis but all genes in this cluster were found in only a subset of all assemblies, the cluster is defined as accessory. Clusters shared by all assemblies are defined as core clusters. If a core cluster had one gene from each assembly, it is defined as ‘strict core’. If one or more assemblies had more than one gene in a core cluster, the cluster is defined as ‘surplus core’.

**Supplementary Fig 2:**
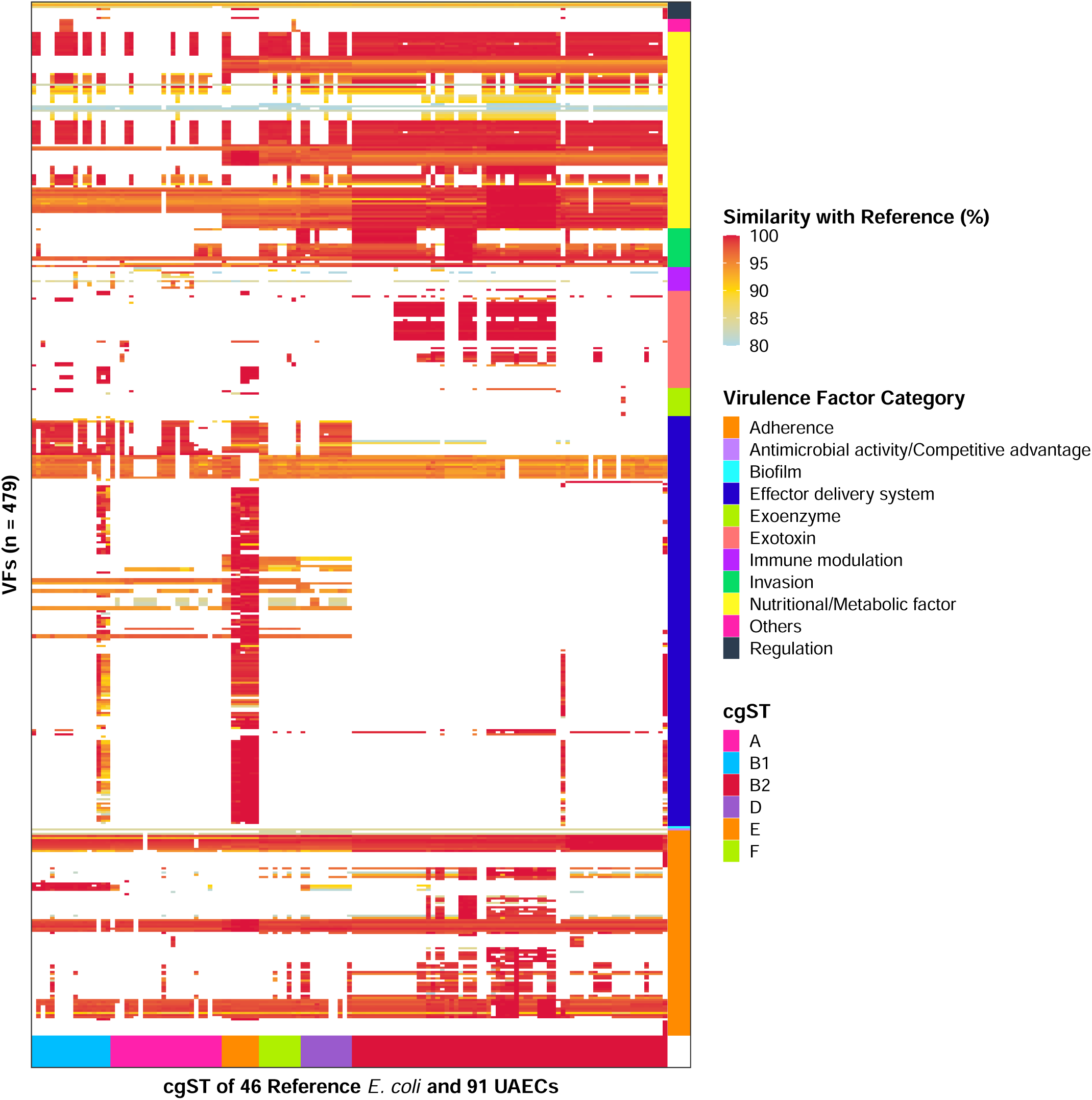
VFs annotation of 46 reference and 91 patient isolated *E. coli*. Plot based on Supplementary Data 1. A fragment owned by strain with similarity against reference lower than 80% was regarded as absence of the corresponding gene in the strain.

**Supplementary Fig 3:**
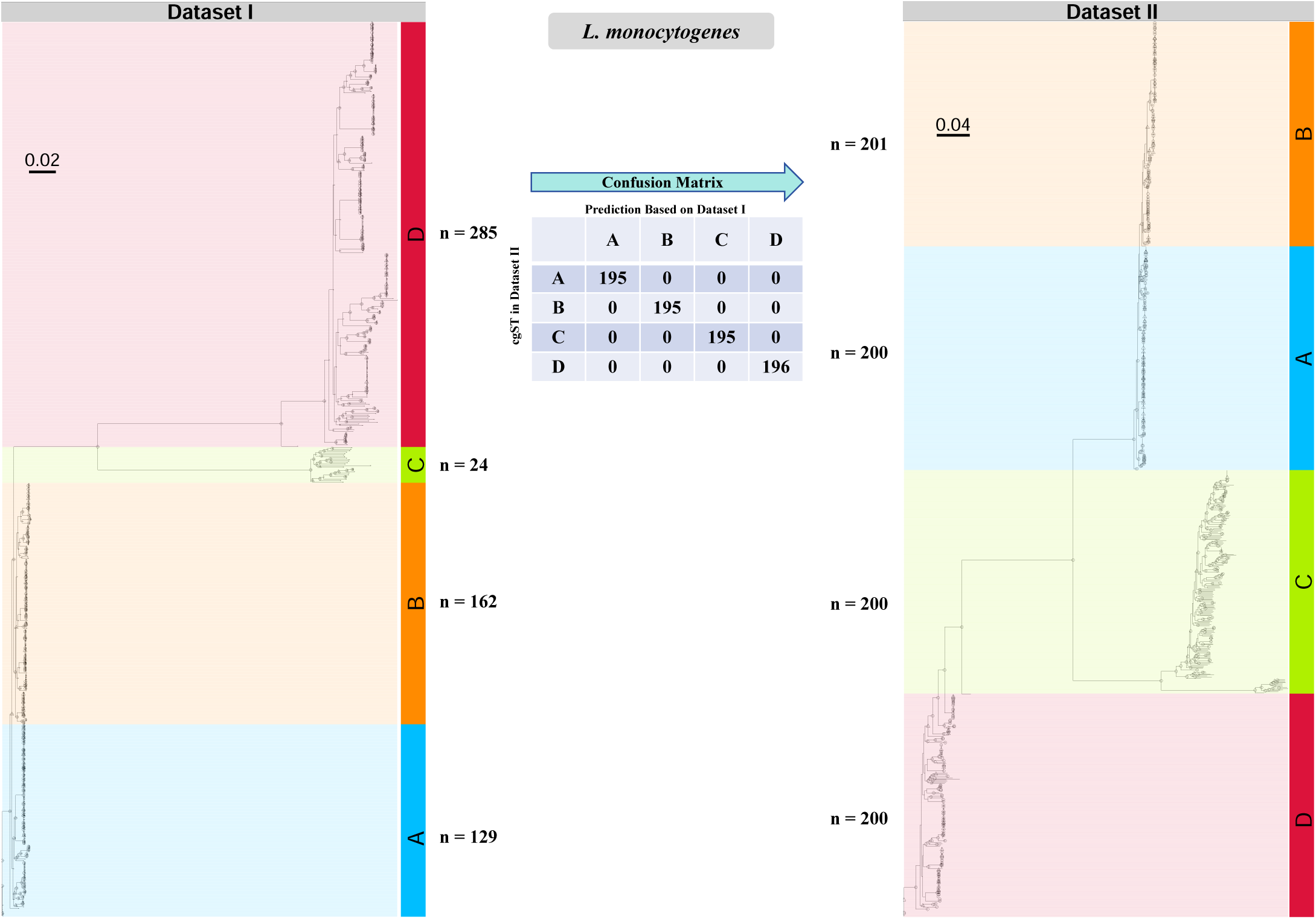
Prediction of cgST by accessory gene status in *L. monocytogenes*. cgST result of Dataset I (left) and II (right) of *L. monocytogenes*. The dashed branches were of large length and were manually shortened to help visualization. The nodes with bootstrap value ≥ 95 are marked with circles and those with bootstrap value >80 and <95 were marked with triangles. The tables in the middle described the confusion matrix which did not include the reference strains (known cgST in Dataset I) and cgST distribution of Dataset I and II next to each cgST.

**Supplementary Fig 4:**
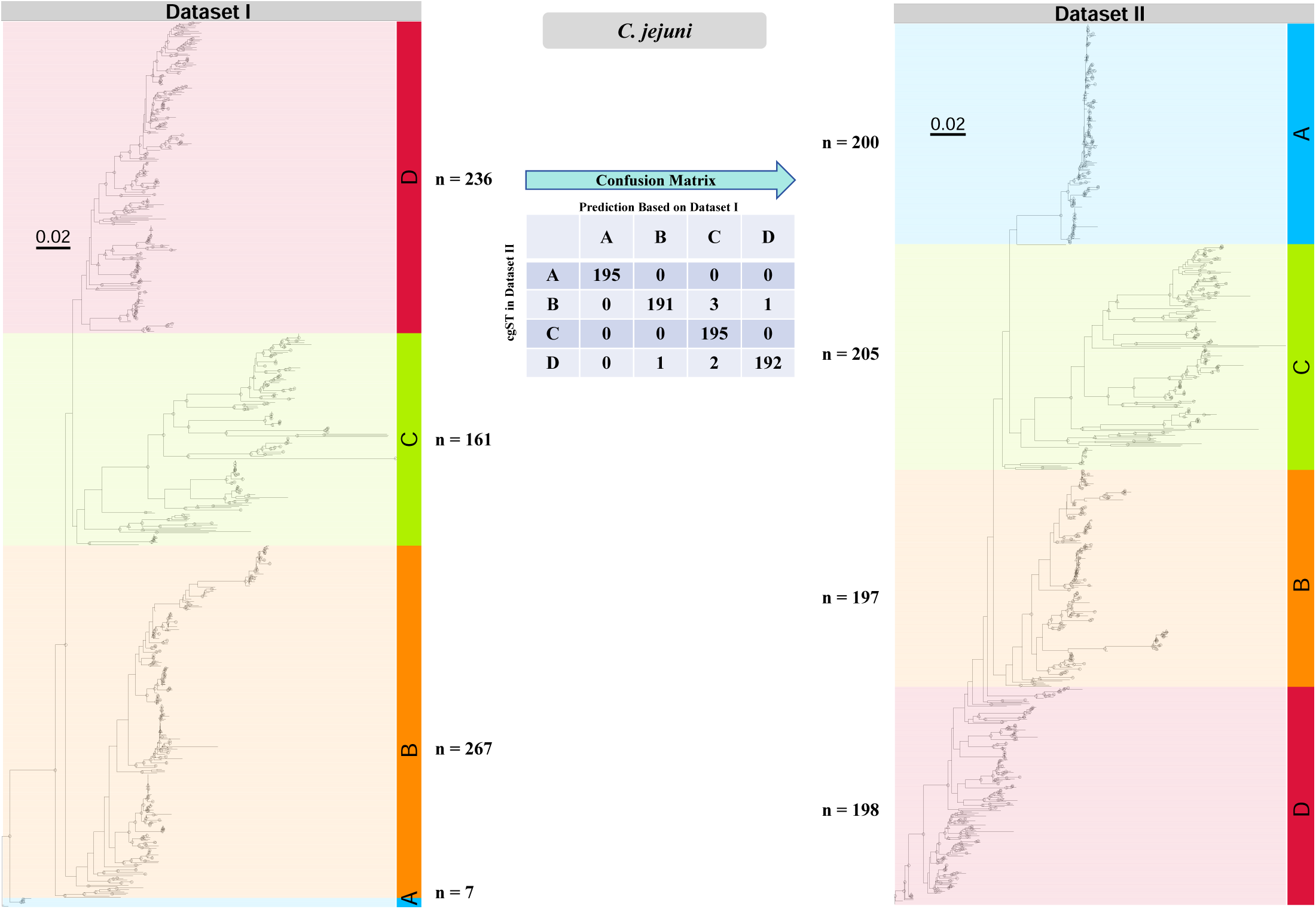
Prediction of cgST by accessory gene status in *C. jejuni*. cgST result of Dataset I (left) and II (right) of *C. jejuni*. The dashed branches were of large length and were manually shortened to help visualization. The nodes with bootstrap value ≥ 95 are marked with circles and those with bootstrap value >80 and <95 were marked with triangles. The tables in the middle described the confusion matrix which did not include the reference strains (known cgST in Dataset I) and cgST distribution of Dataset I and II next to each cgST.

**Supplementary Fig 5:**
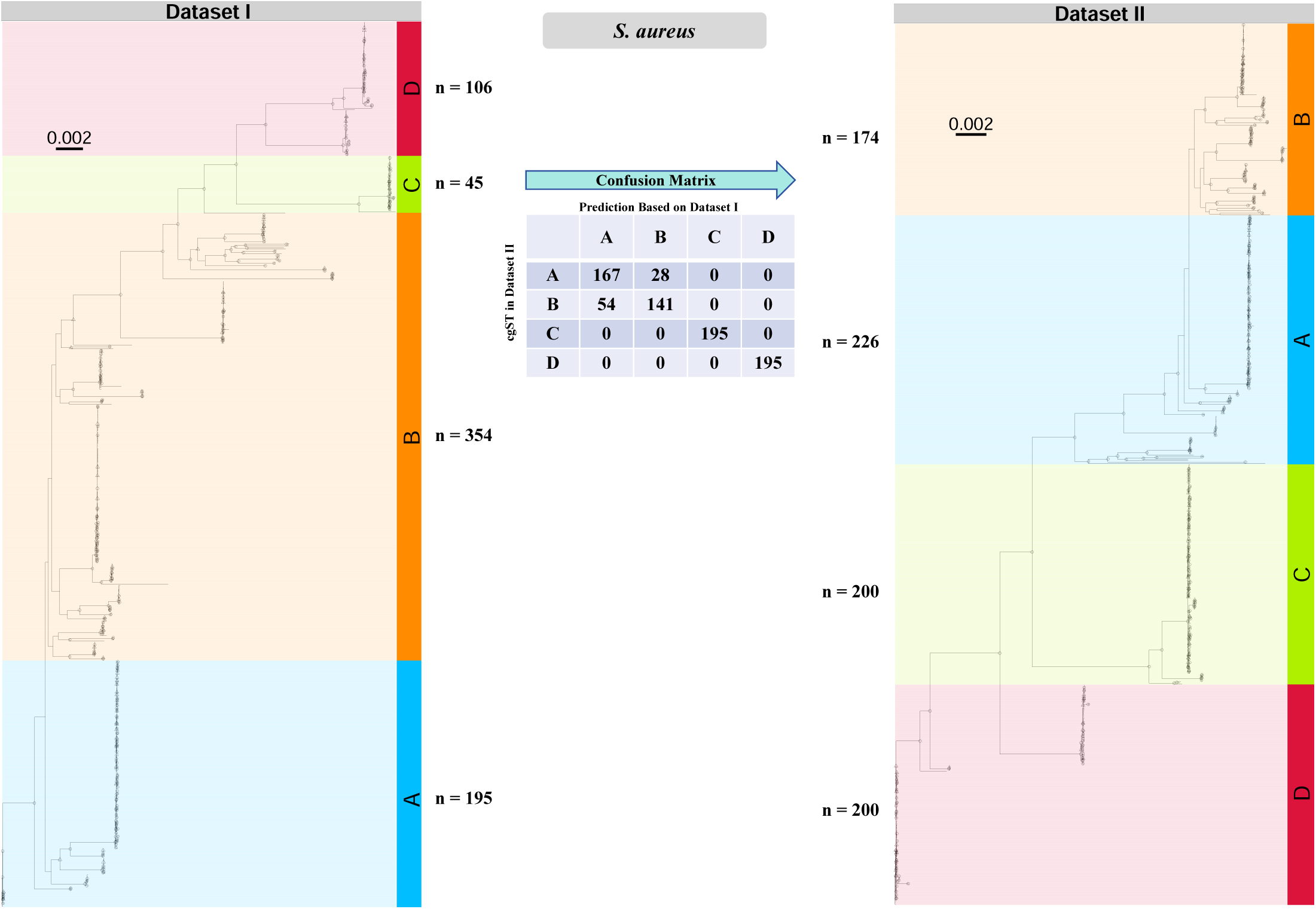
Prediction of cgST by accessory gene status in *S. aureus*. cgST result of Dataset I (left) and II (right) of *S. aureus*. The dashed branches were of large length and were manually shortened to help visualization. The nodes with bootstrap value ≥ 95 are marked with circles and those with bootstrap value >80 and <95 were marked with triangles. The tables in the middle described the confusion matrix which did not include the reference strains (known cgST in Dataset I) and cgST distribution of Dataset I and II next to each cgST.

**Supplementary Fig 6:**
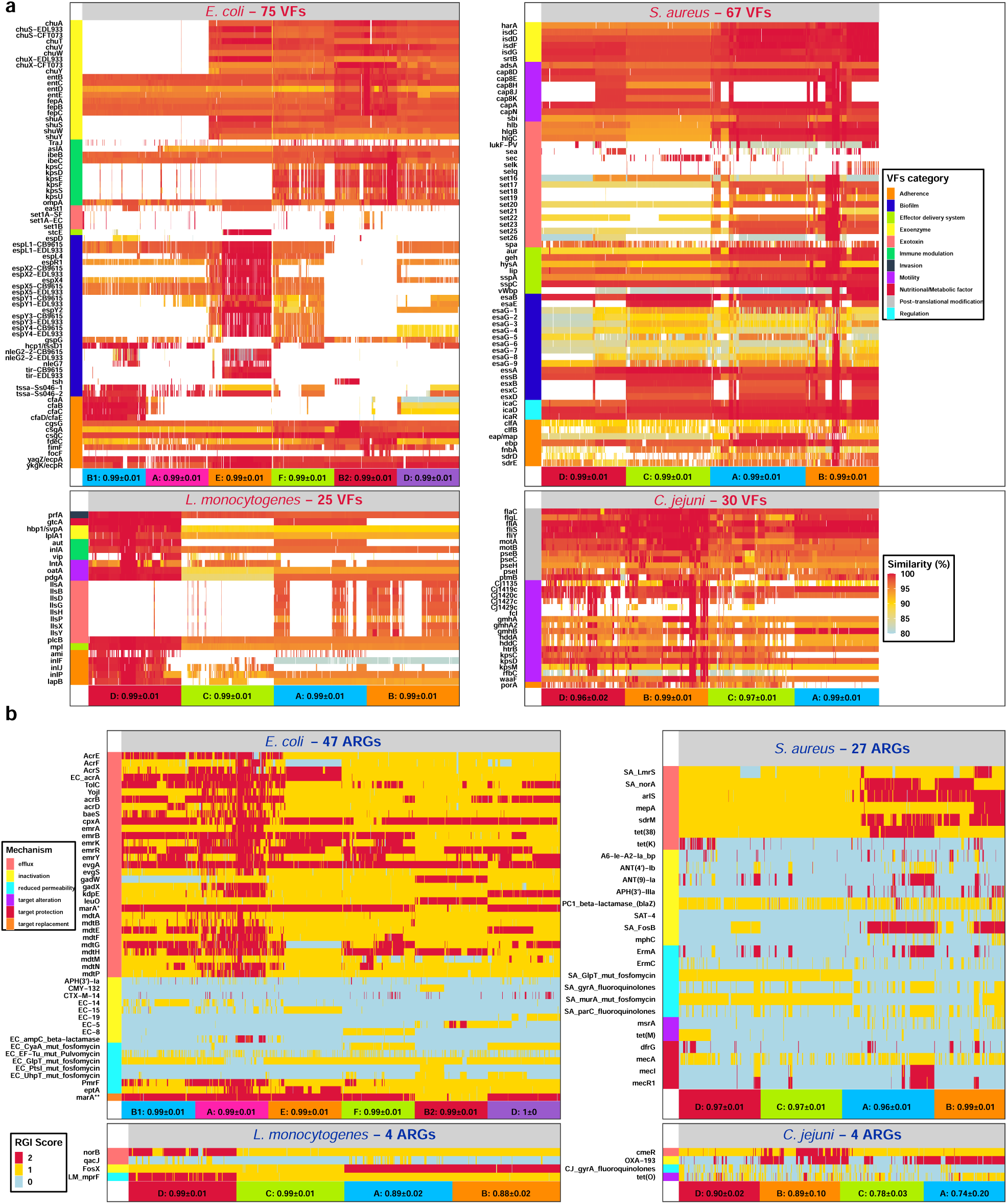
VFs (a) and ARGs (b) specific to cgST were selected by LR based re-trains. The x axis of each plot are strains in Dataset II and are arranged in the same order as in the phylogeny of Dataset II (Fig 2 and Supplementary Fig 2-4) and with same color. The text on x axis also exhibited the average AUROC of re-trains when using VFs (**a**) or ARGs (**b**) patterns to predict cgST. The color on y axis suggests the VFs category (**a**) and resistance mechanism (B) of genes. VFs (**a**) and its VFDB query correspondence could be found in Supplementary Data 6. ARGs (**b**) with simplifies model names correspond to their full name in CARD database in Supplementary Data 6. In plot B, *marA* with “*” suggests the gene was present in 2 different resistance mechanisms.

**Supplementary Fig 7:**
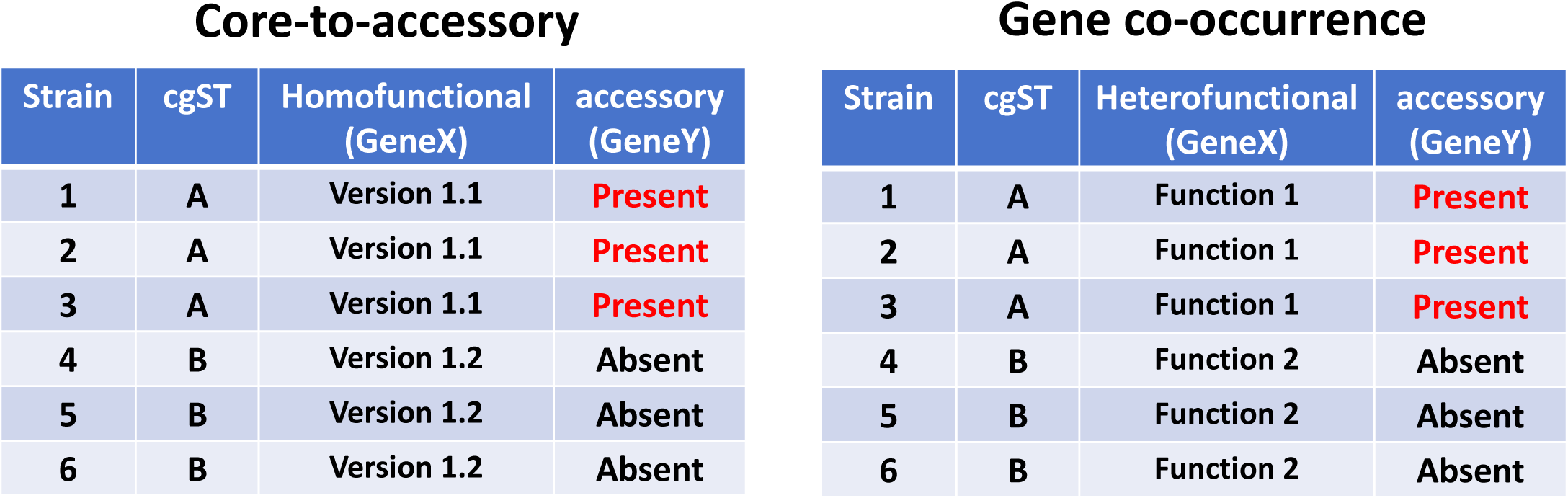
Diagram of difference between core-to-accessory relationship and gene co-occurrence. In this diagram, we show two possible modes that accessory genes can correlate with certain cgST. In this simplified example, 6 strains were classified into 2 different cgSTs (A and B) by a singe hypothetical core gene cluster (name as GeneX) shared by all 6 strains. However, GeneX from six strians might be of same function (left table), where version 1.1 of GeneX might have better efficiency than version 1.2. The GeneX from cgST A and B might also come from different function (right table) due to misclustering therefore we mistakenly regarded two accessory clusters as a core cluster. In this case, the specific presence of GeneY in cgST A, is a co-occurrence with GeneX of Function 1.

**Supplementary Fig 8:**
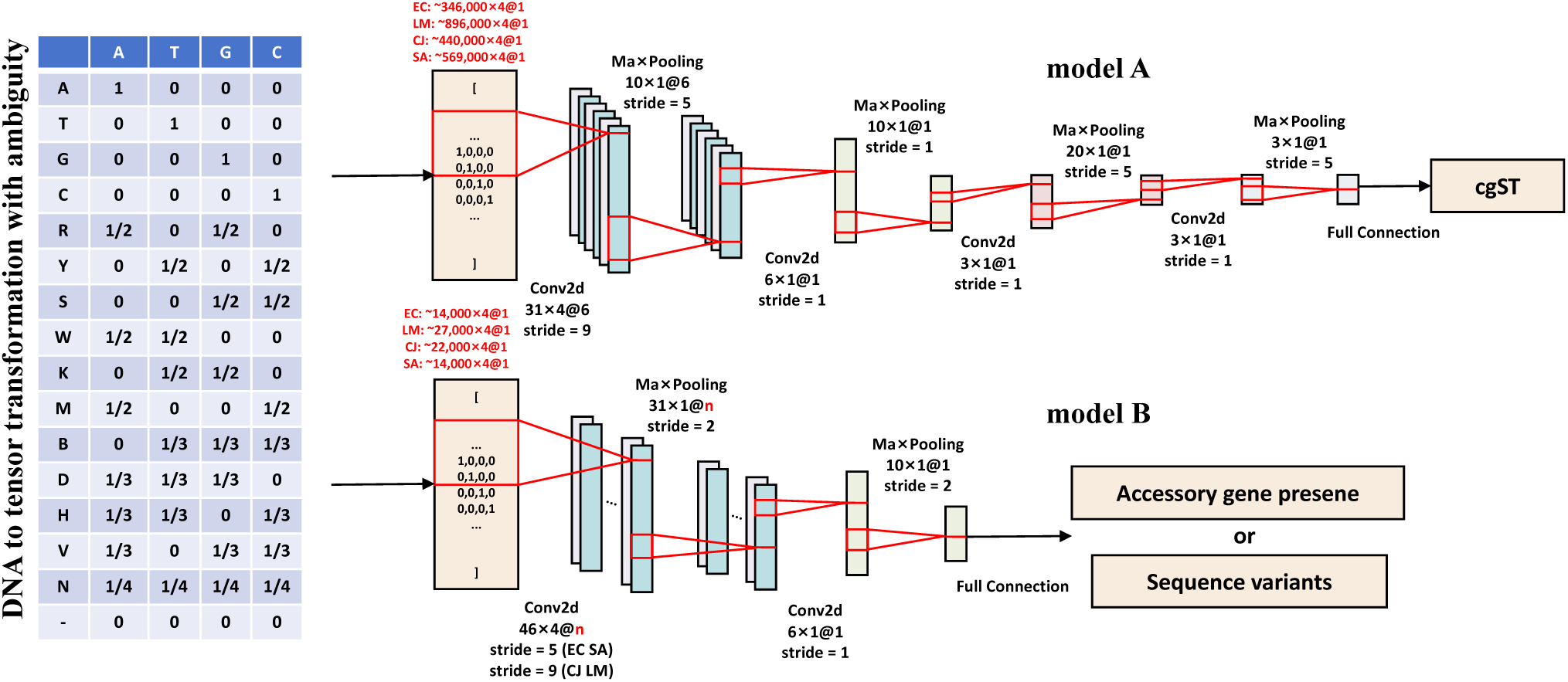
Transform DNA to tensor using one-hot like encoding and applying predictions with CNN model A and B. Input DNA sequences were first transformed using a one-hot like manner. The transformation put IPUAC DNA ambiguity into considerations. Model A were using for cgST prediction which included 4 convolution layers and 4 maxpooling layers and only the first maxpooling and convolution layers had 6 channels. Model had a deeper structure than model B due input size. Model A took complete core genome size as input (∼346,000 bp to 890,000 bp), therefore, a deeper structure for feature extraction and dimension reduction. While model B took partial genome (∼14,000 bp to 27,000 bp) as input to predict accessory gene presence and sequence variant of other genes. Model B had only 2 convolution layers and 2 maxpooling layers and the amount of channels for first convolution and maxpooling layers was equal to the count of class (n = count of sequence variants) to be predicted. The channel count in model B was set to 6 in phenotype prediction.

**Supplementary Fig 9:**
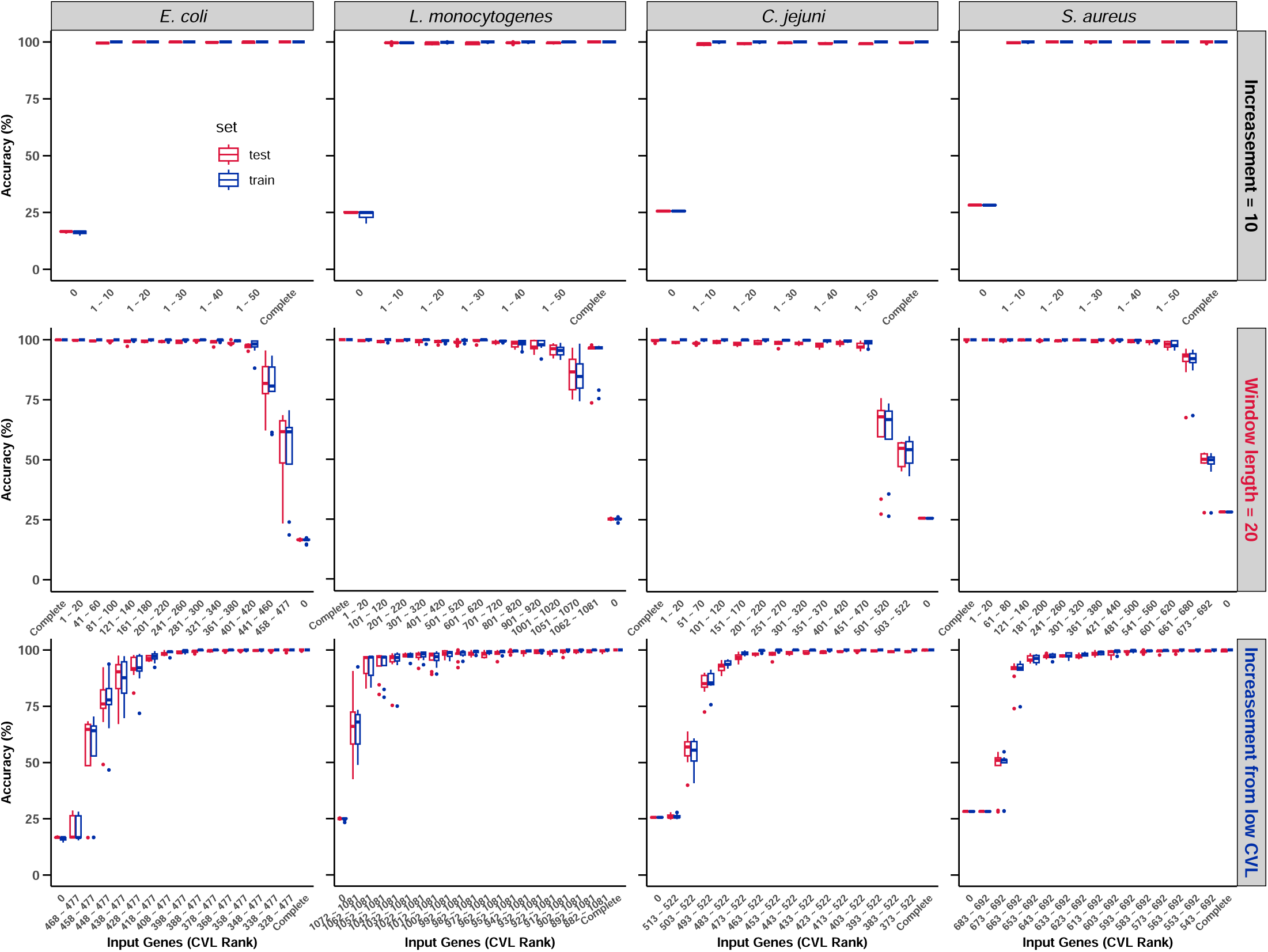
Different core gene combinations can accurately recover cgSTs with CNN in cross validations. (EC: *E. coli*, LM: *L. monocytogenes*, CJ: *C. jejuni*, SA: *S. aureus*). First row: by increasing available genes in the input with increase of ten genes (from the high CVL StCGCs), cgST recovery of >98% could be achieved with less than 20 StCGCs. Second row: By using different window (20 genes each window) of available core genes in prediction, ability to recover complete core genome cgST was tested on a wider range of core genome. Third row: by increasing available genes from the lower end from the CVL list, least needed number to recover cgST with >98% accuracy from the bottom of CVL list is exhibited. Each box represents the performance of nine re-trains (three-fold, three random initialization each fold) of train set (red) and test set (blue). The CNN model adopted model A as in Supplementary Fig 8.

**Supplementary Fig 10:**
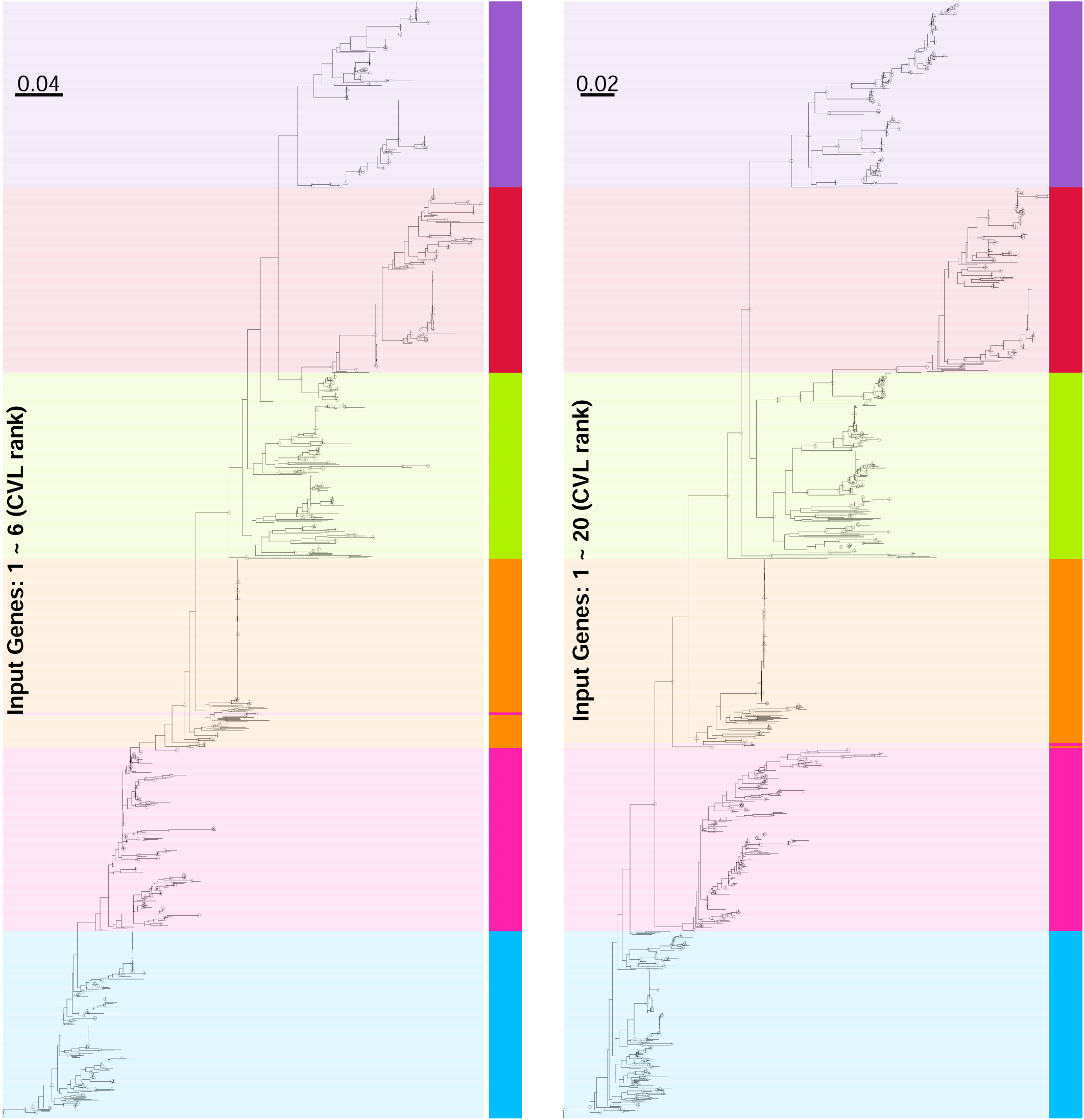
*E. coli* Dataset II phylogeny constructed using subset of StCGCs. Left tree used gene 1 ∼ 6 and right tree used gene 1 ∼ 20 in CVL list. Dashed branch was a outlier with too large branch length manually shortened to help visualization. The nodes with bootstrap value ≥ 95 are marked with circles and those with bootstrap value >80 and <95 were marked with triangles. The color for each strain is the same as in Fig 2.

**Supplementary Fig 11:**
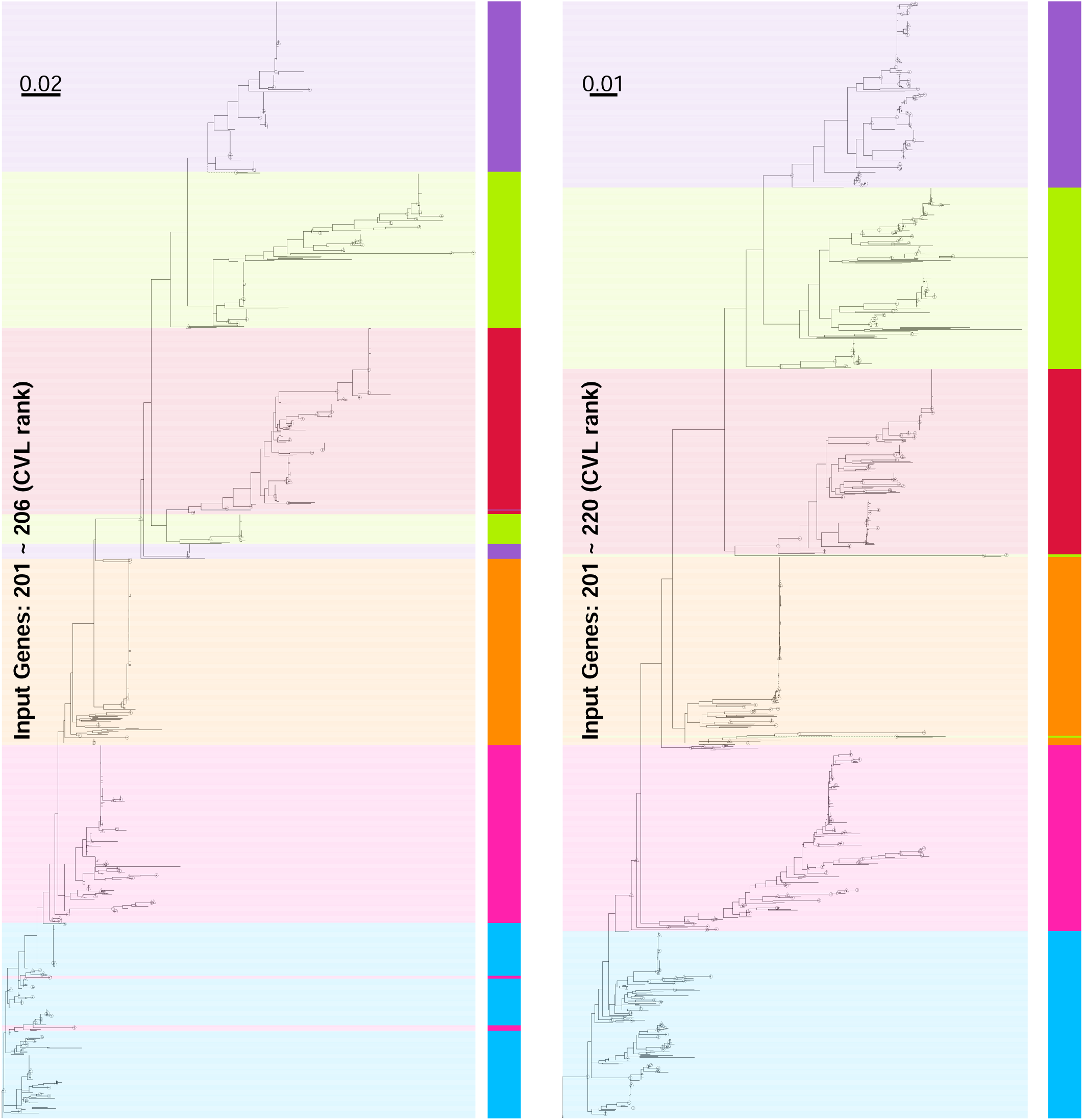
*E. coli* phylogeny constructed using subset of StCGCs. Left tree used gene 201 ∼ 206 and right tree used gene 201 ∼ 220 in CVL list. Dashed branch was a outlier with too large branch length manually shortened to help visualization. The nodes with bootstrap value ≥ 95 are marked with circles and those with bootstrap value >80 and <95 were marked with triangles. The color for each strain is the same as in Fig 2.

**Supplementary Fig 12:**
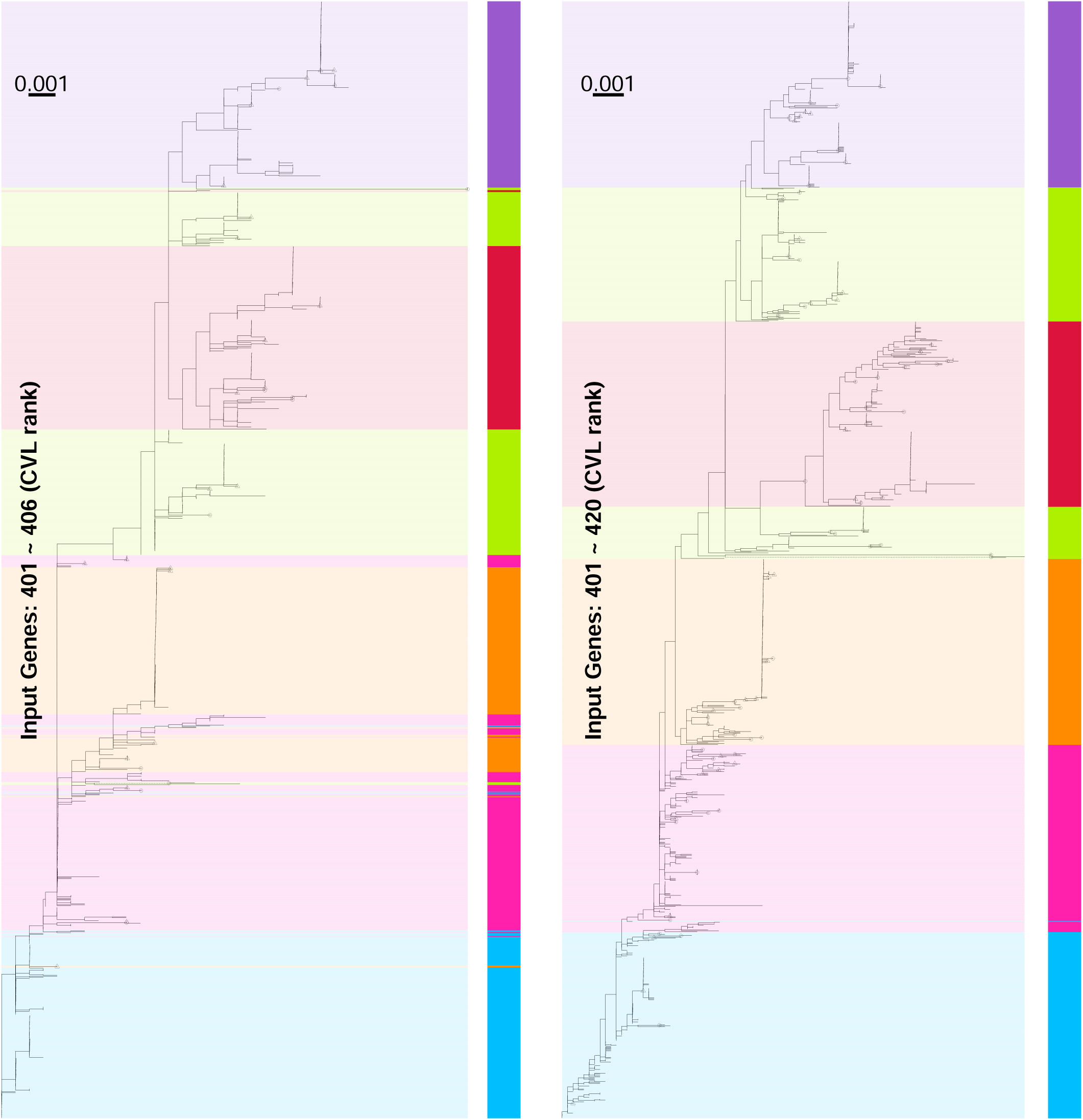
*E. coli* phylogeny constructed using subset of StCGCs. Left tree used gene 401 ∼ 406 and right tree used gene 401 ∼ 420 in CVL list. Dashed branch was a outlier with too large branch length manually shortened to help visualization. The nodes with bootstrap value ≥ 95 are marked with circles and those with bootstrap value >80 and <95 were marked with triangles. The color for each strain is the same as in Fig 2.

**Supplementary Fig 13:**
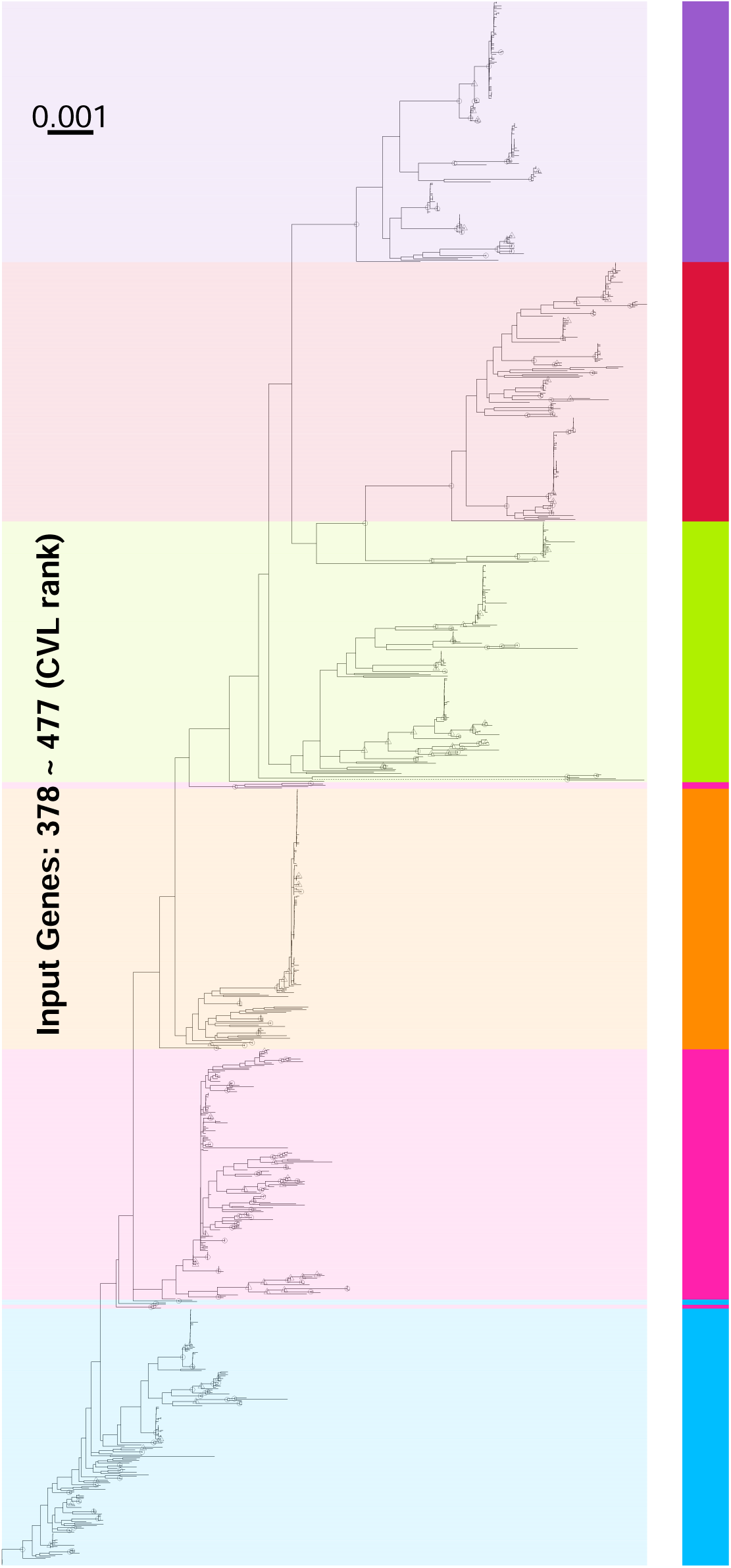
*E. coli* phylogeny constructed using subset of StCGCs. The phylogeny was constructed using last 100 genes (378 ∼ 477) in CVL list. Dashed branch was a outlier with too large branch length manually shortened to help visualization. The nodes with bootstrap value ≥ 95 are marked with circles and those with bootstrap value >80 and <95 were marked with triangles. The color for each strain is the same as in Fig 2.

**Supplementary Fig 14:**
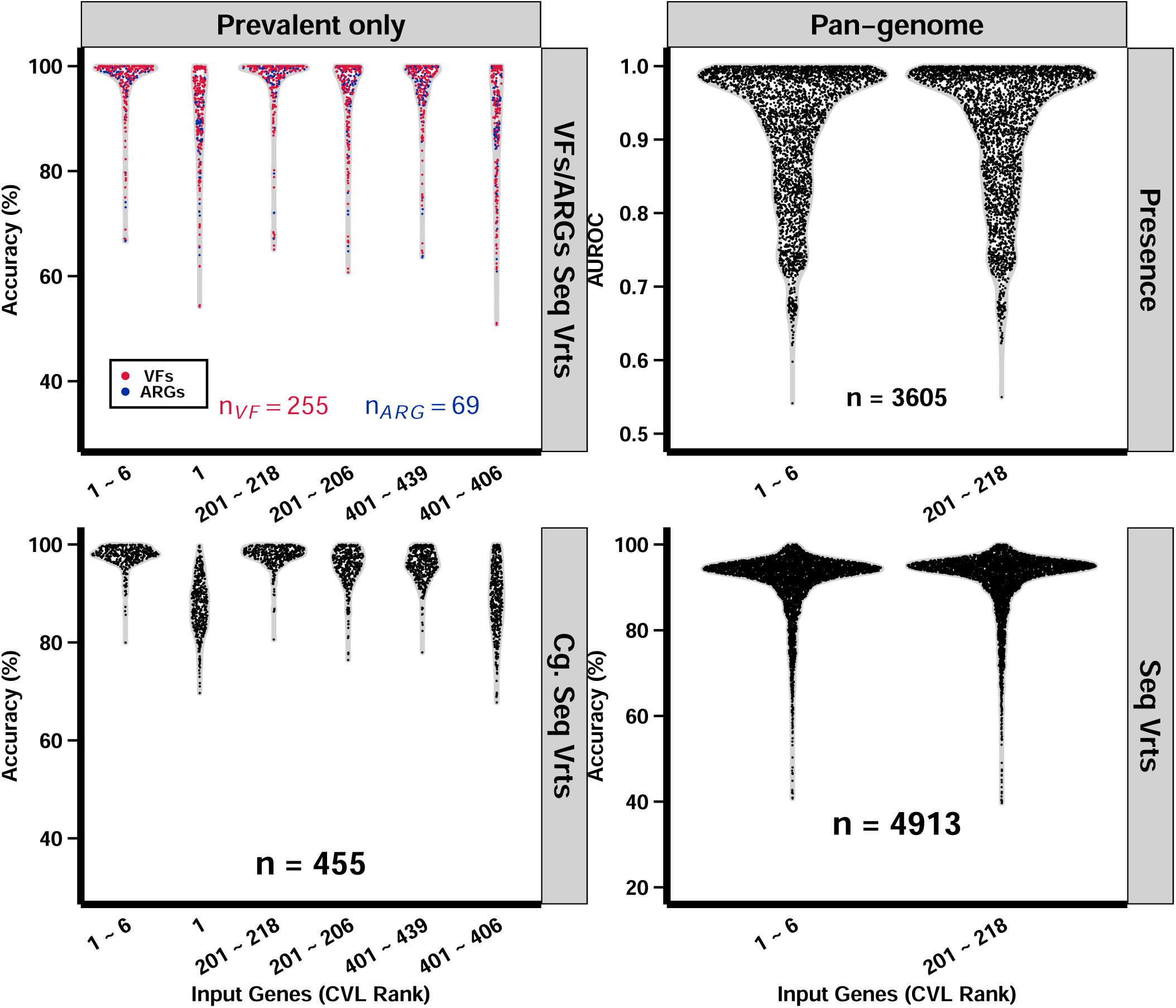
Predict gene status using limited core genes, embedded core genes, and correlation between frequency and predictability. **First column**: the prediction of sequence variant (Seq Vrts) of VFS/ARGs and core genes (Cg.) were tested when only the prevalent sequence variants were predicted (excluding the class ‘other’). **Second column**: with embedded input, the gene status prediction of a large range of genes from the pan-genome was tested and achieved a high average predictability. **Third column**: predictability of genes in the pan-genome was test for correlation with genes frequency (presence rate) in SAC in which gene status predictability do not show correlation with gene frequency.

**Supplementary Fig 15:**
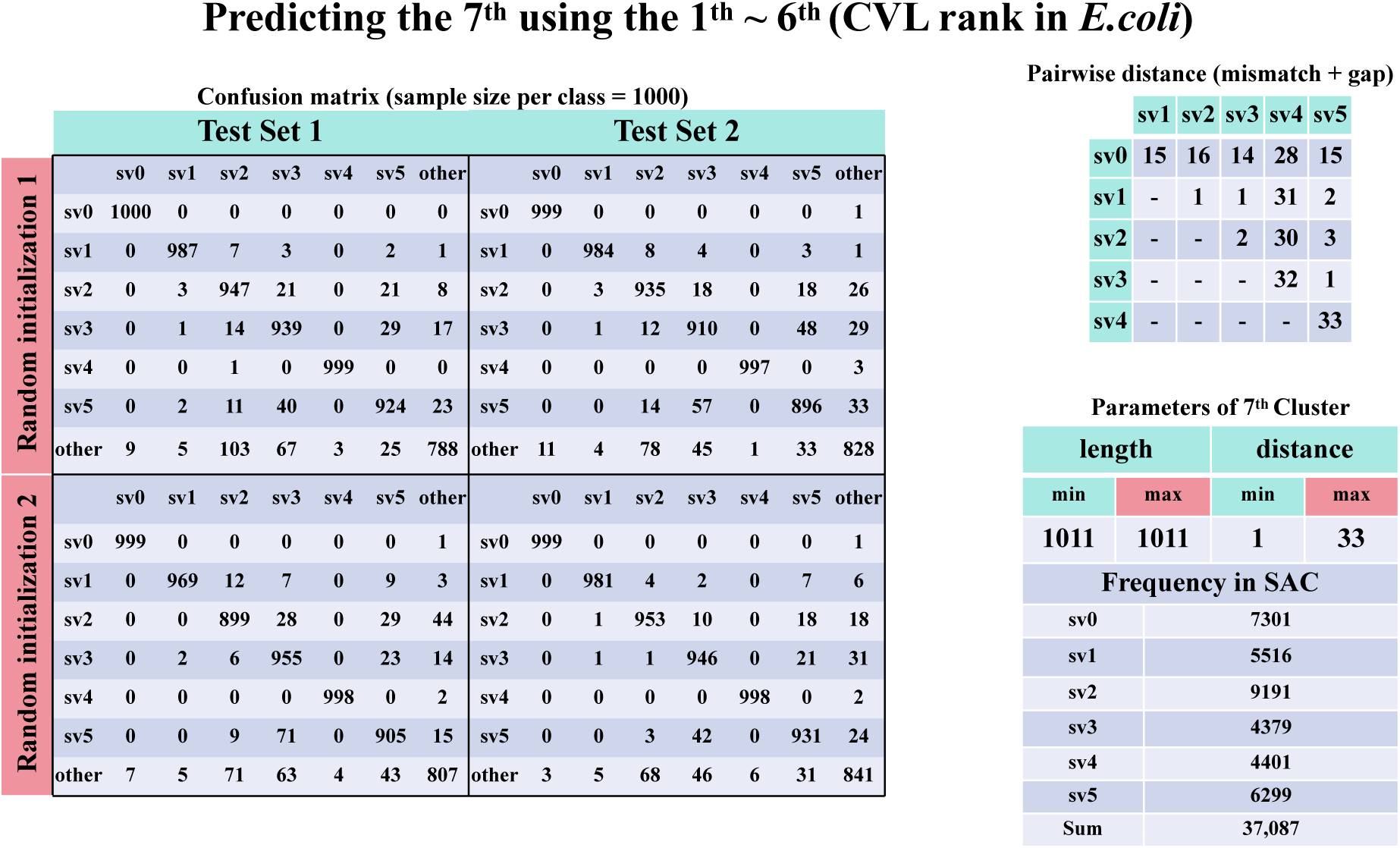
Results of predicting *E.coli* 7^th^ (CVL rank) gene using 1^st^ to 6^th^ and different between sequence variants. The confusion matrix of each four re-trains (**left**), two random initialization within each of two-fold rows, had rows as actual class and columns as predicted class where each class has 1,000 samples. The table on **top right** indicated the pariwise distance (mismatches + gaps) among all prevalent sequence variants (sv). In **bottom right**, the table indicated that all prevalent st has same length therefore the minimum (min) distance is 1 nucleotide difference over all pairwise distances; this table also indicated the frequency of each prevalent st in the SAC.

**Supplementary Fig 16:**
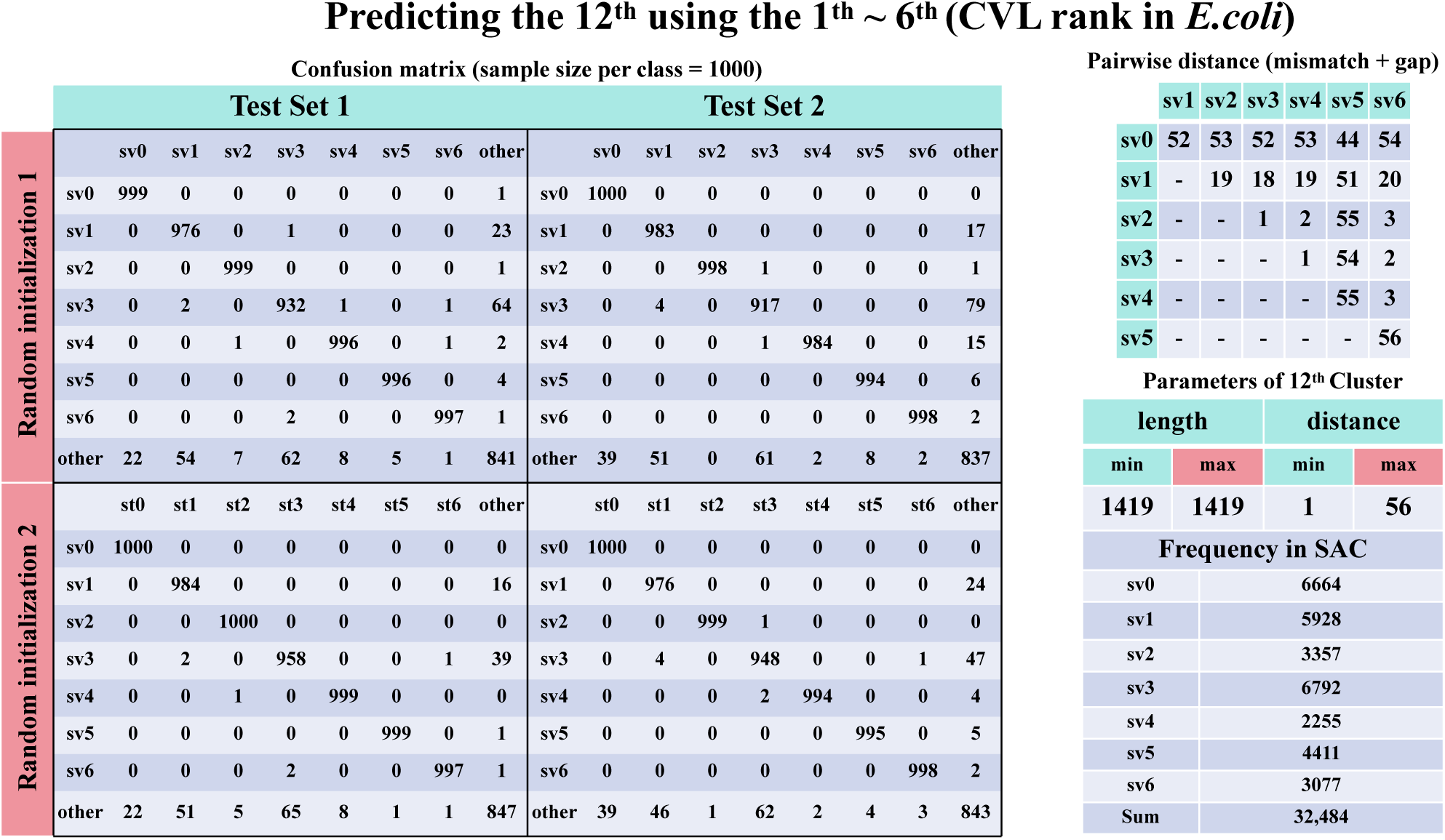
Results in predicting *E.coli* 12^th^ (CVL rank) gene using 1^st^ to 6^th^ and different between sequence variants. The confusion matrix of each four re-trains (**left**), two random intialization within each of two-fold rows, had rows as acutal class and columns as predicted class where each class has 1,000 samples. The table on **top right** indicated the pariwise distance (mismatches + gaps) among all prevalent sequence variants (sv). In **bottom right**, the table indicated that all prevalent st has same length therefore the minimum (min) distance is 1 nucleotide difference over all pairwise distances; this table also indicated the frequency of each prevalent st in the SAC.

**Supplementary Fig 17:**
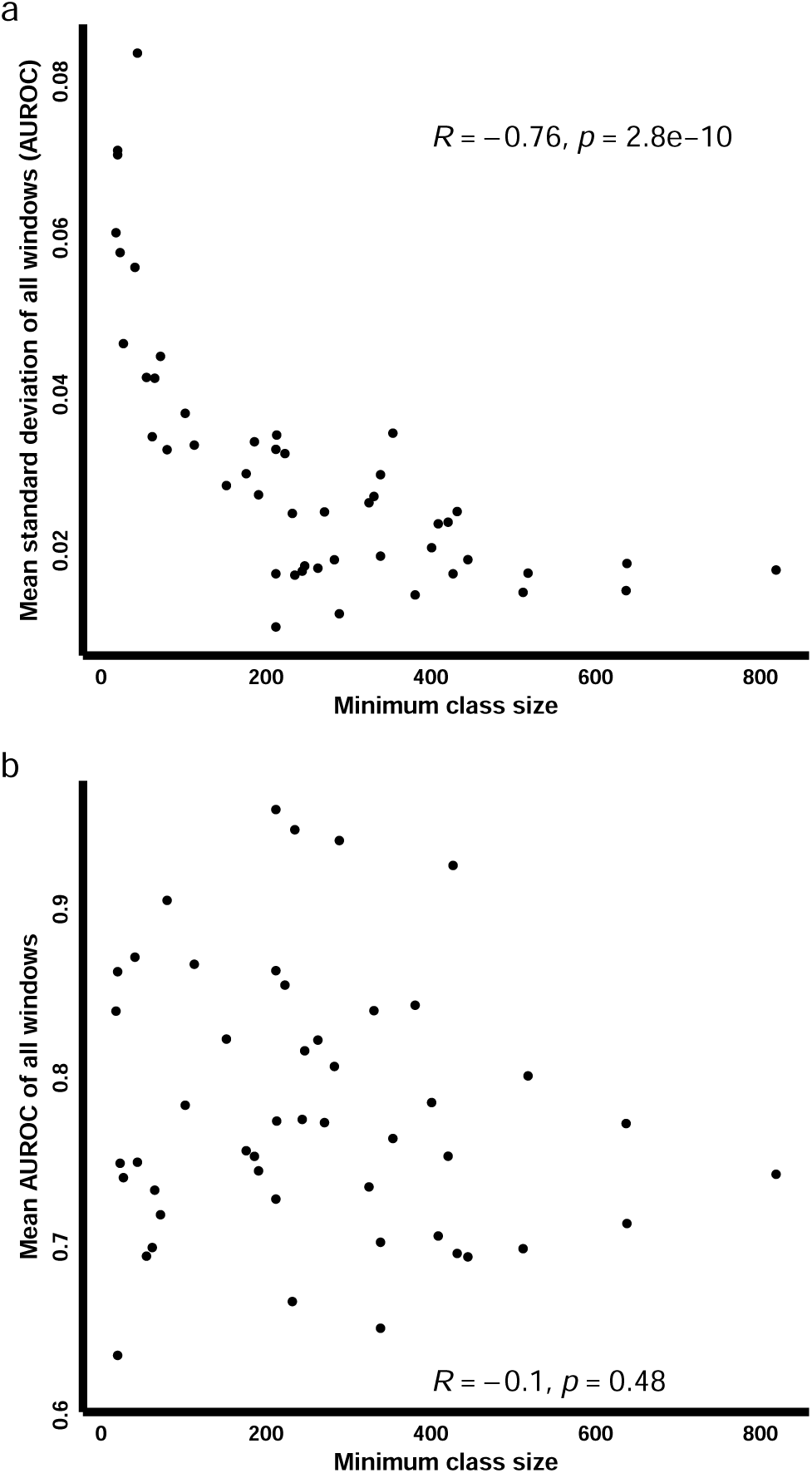
Sample size available in training correlates with consistent test set cross validation performance (a) but not with better prediction accurary (b). x axis of both **a** and **b** is the minimum class size, as the minimum number among resistant and susceptible sample size (Fig 5). **(a)** y axis represents the mean standard deviation of all input windows of a phenotype prediction. **(b)** y axis represents the mean prediction performance of all input windows of a phenotype prediction.

**Supplementary Fig 18:**
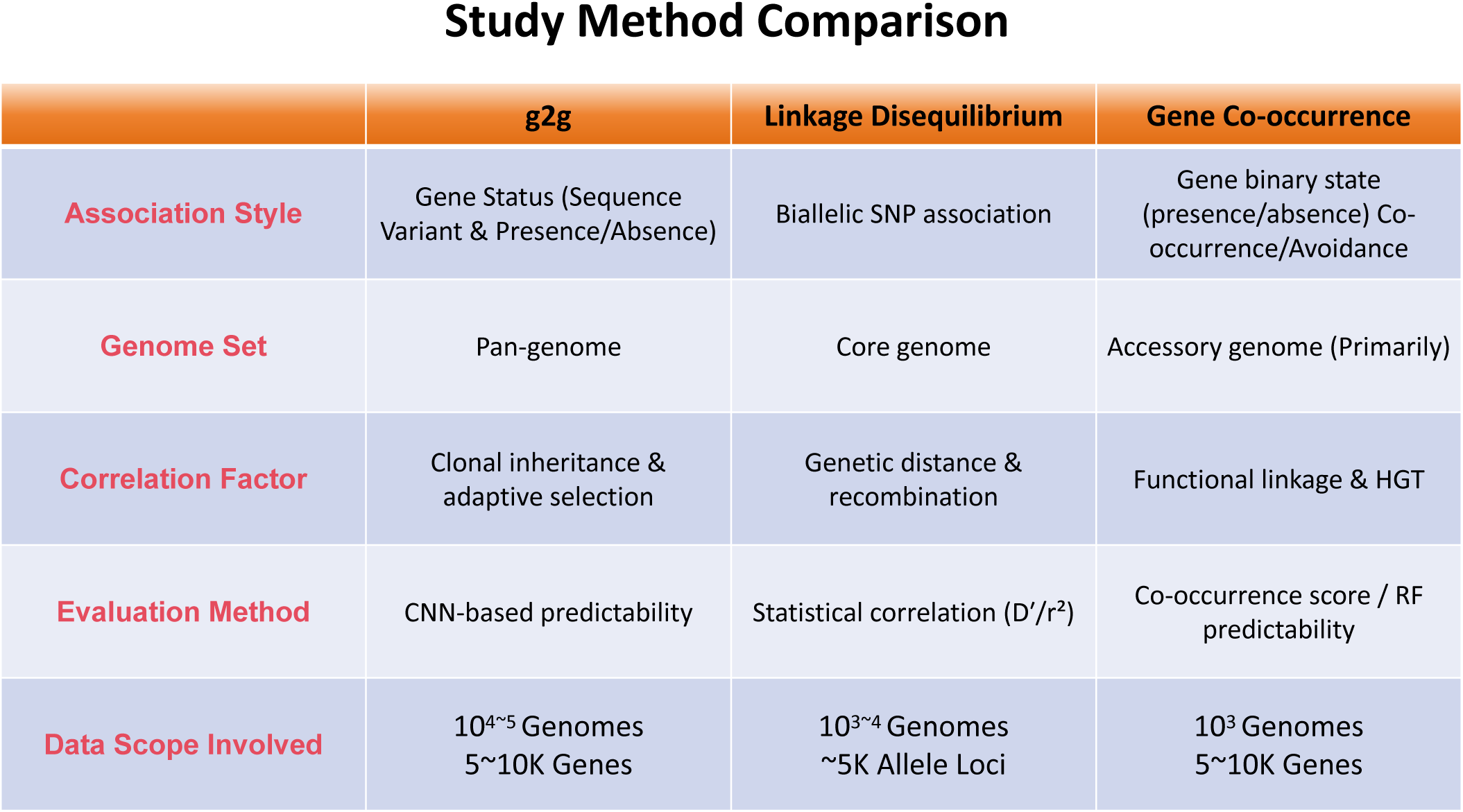
Study method comparison among g2g, Linkage Disequilibrium, and Co-occurrence. The g2g correlation method presented in this study, classic Linkage Disequilibrium (LD), and Gene Co-occurrence analysis. The g2g approach is distinct in its use of multi-state gene status (integrating both sequence variants and presence/absence) across the pan-genome, evaluated through CNN-based predictability. This allows it to capture complex correlations driven by clonal inheritance and adaptive selection on a large scale (10 –10 genomes, 5-10k genes). In contrast, LD typically analyzes biallelic SNP associations in the core genome, measuring statistical correlations influenced by genetic distance and recombination. Gene Co-occurrence analysis focuses on presence/absence patterns in the accessory genome, identifying functional linkages through statistical scores or Random Forest models. The g2g framework provides a more comprehensive and high-resolution view of genomic correlations, overcoming the limitations of binary-state analyses.

**Supplementary Fig 19:**
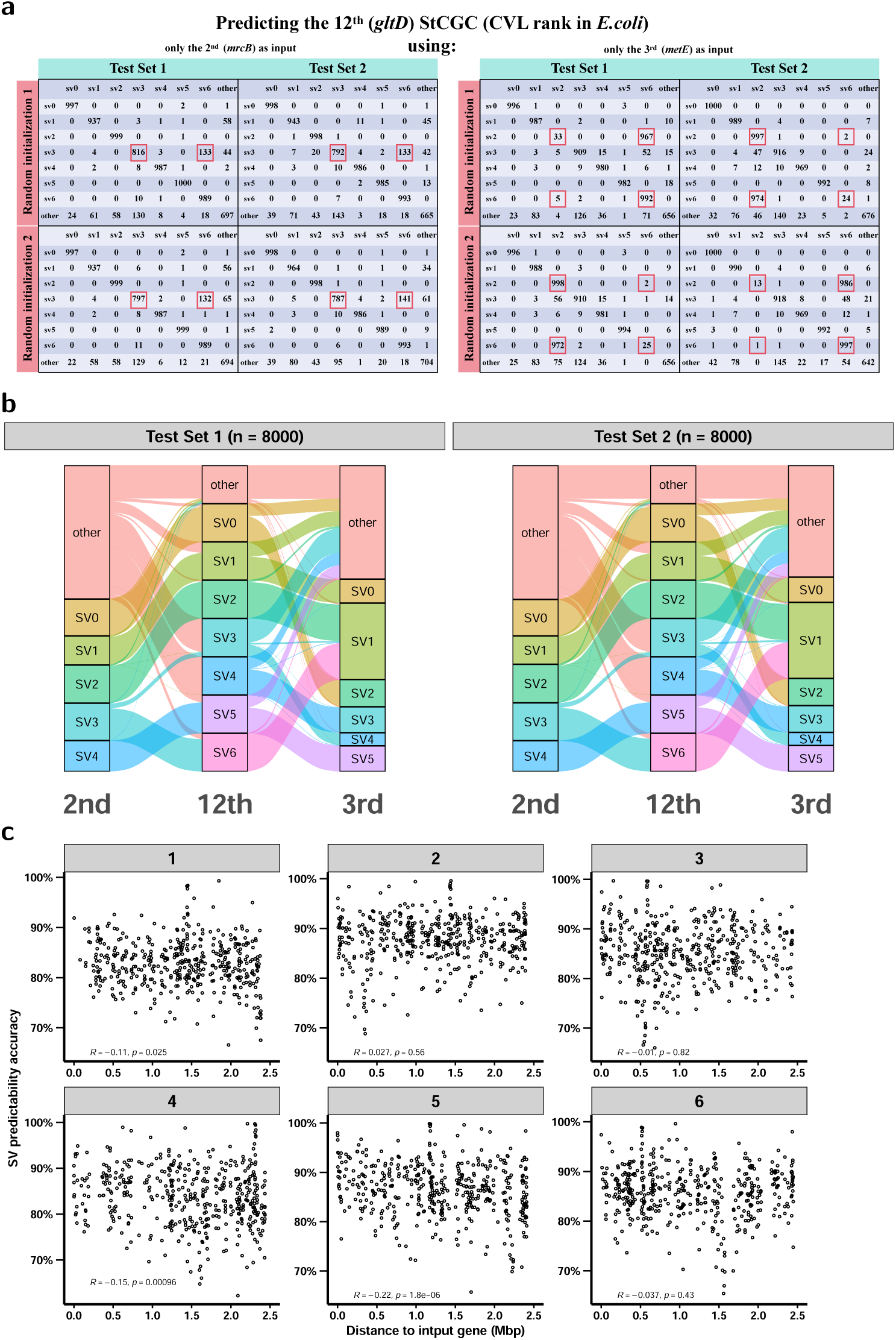
Association complexity of g2g. **(a)** Confusion matrices showing prediction performance for *gltD* SVs using only the 2^nd^ CVL-ranked gene (*mrcB*) as input (left) or only the 3^rd^ CVL-ranked gene (*metE*) as input (right). Test set 1 and 2 are non-overlapping collections, with each *gltD* SV having 2,000 strains. Cells with red frames indicate SV pairs prone to misprediction. **(b)** Sankey plots depicting the complexity of SV combinations among the 2^nd^ (*mrcB*), 3^rd^ (*metE*), and 12^th^ (*gltD*) CVL-ranked genes in the same Test Set 1 and 2 as panel **(a)**. Flows represent the distribution of SVs across these genes. **(c)** Grey strip on top of each sub-plot indicates the CLV rank of each single prediction input. Each sub-plot describes the weak correlation (spearman) between SV prediction accuracy over distance (Mbp) to the genes being predicted.

**Supplementary Fig 20:**
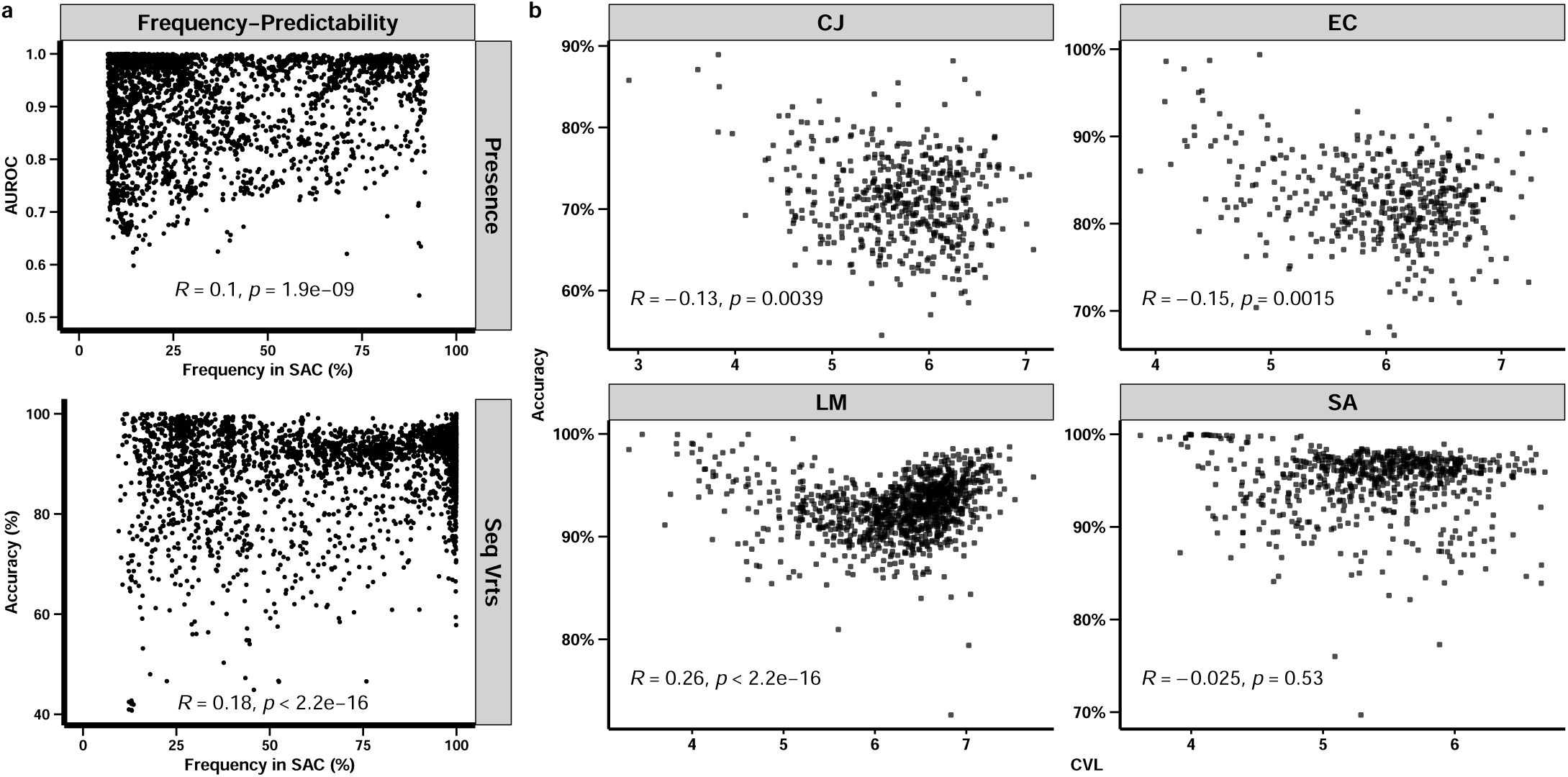
Weak predictability correlation withother factors. **(a)** The weak correlation (spearman) between gene status predictability over frequency in the SAC. **(b)** describes the weak correlation (spearman) between CVL value and predictability (accuracy) of StCGCs of Dataset II of the four species. EC: *E. coli*, LM: *L. monocytogenes*, CJ: *C. jejuni*, SA: *S. aureus*.

**Supplementary Fig 21:**
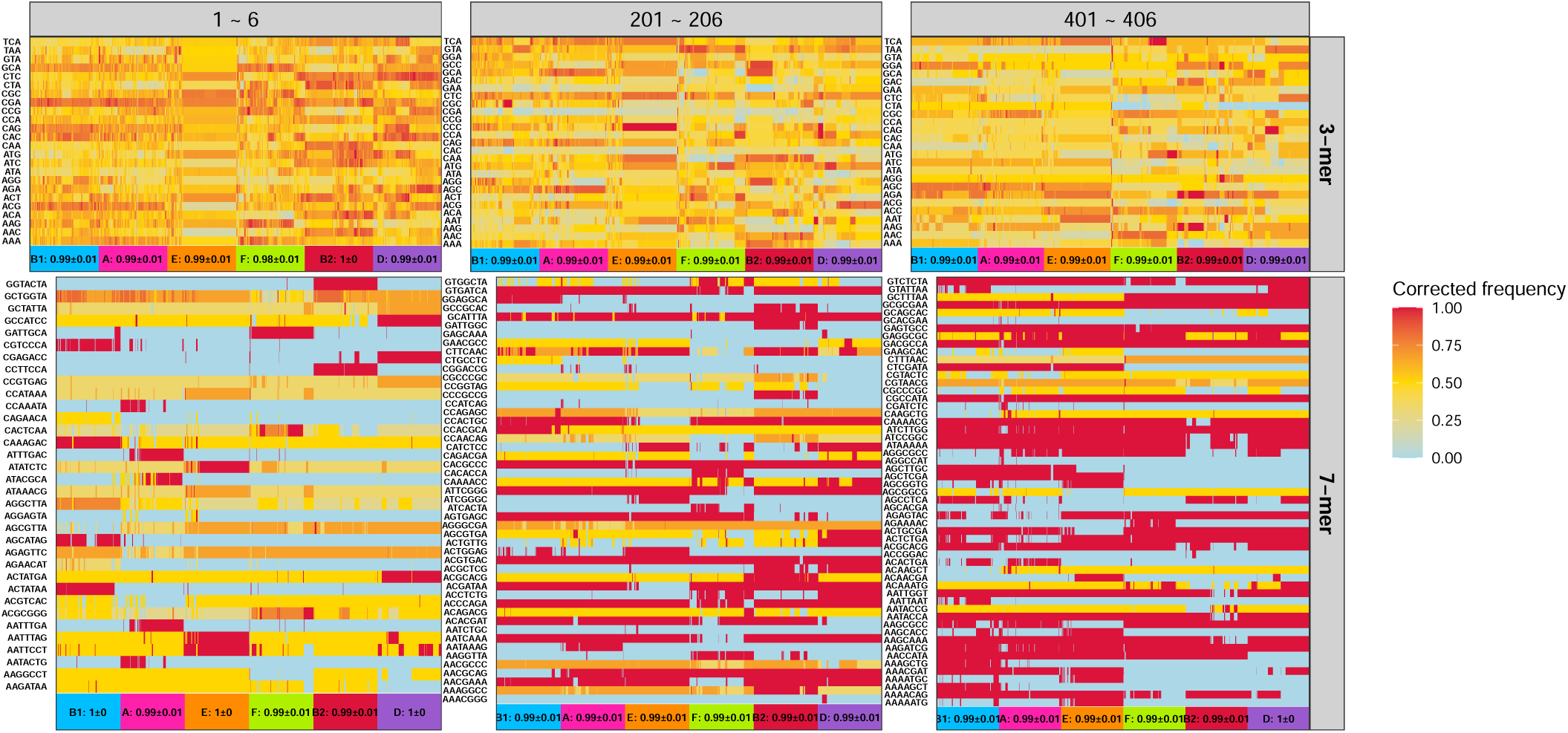
*E. coli* cgST specific k-mers selected by LR based re-trains. 3-mer (first row) and 7-mer (second row) frequency were extracted from Dataset II of *E. coli*. These k-mer patterns from StCGCs CVL rank window of 1 to 6 (first column), 201 to 206 (second column), and 401 to 406 (third column) were used to predict stain’s cgST where re-train were performed 20 times for each cgST. Strains on the x axis were stained the same color as in Fig 2 with cgST corresponding color. Number on the x axis is the average AUROC over 20 re-trains in predicting the cgST.

**Supplementary Fig 22:**
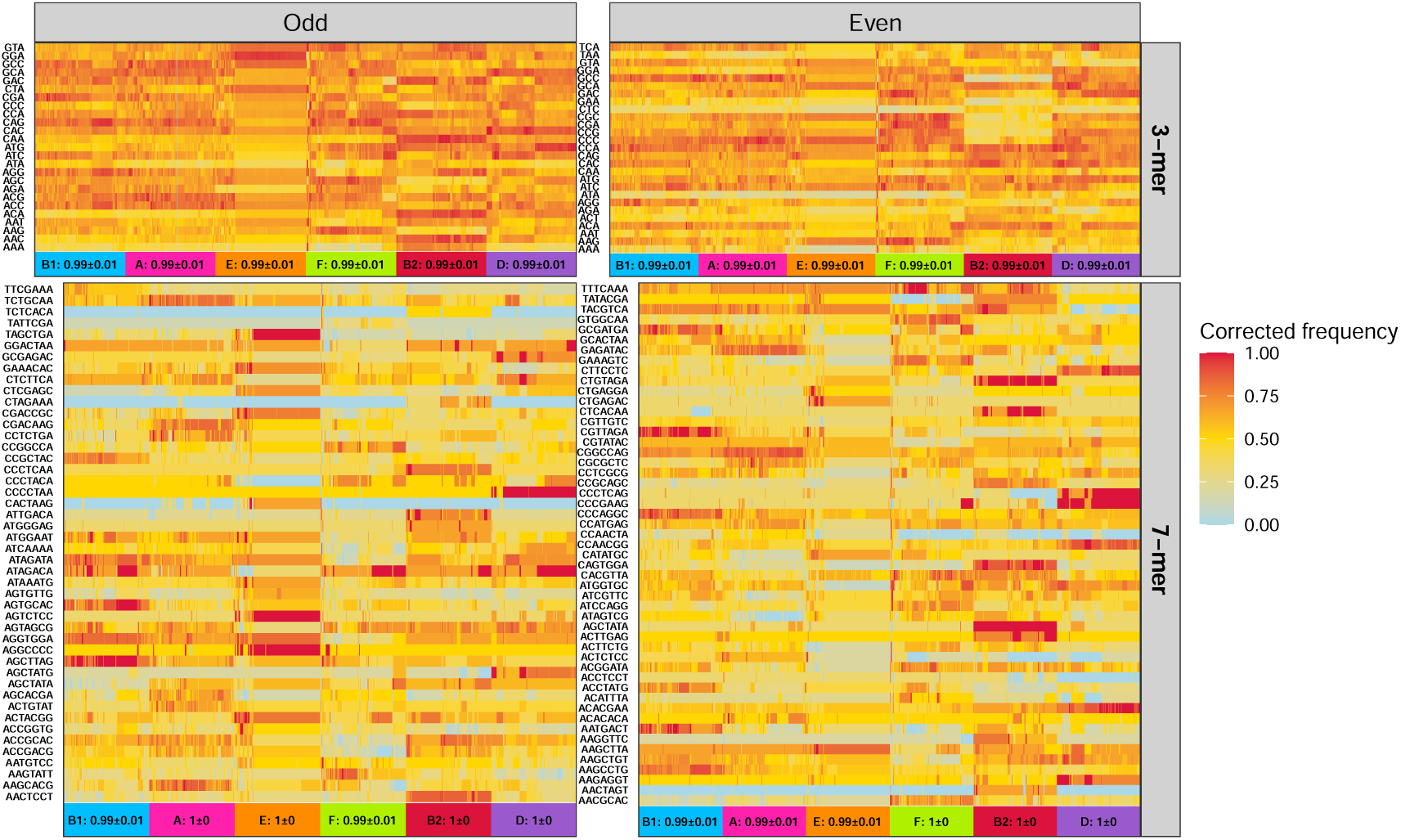
*E. coli* cgST specific k-mers selected by LR based re-trains. 3-mer (first row) and 7-mer (second row) frequency were extracted from Dataset II of *E. coli*. And these k-mer patterns from StCGCs rank list with odd (first column) and even (second column) rank were used to predict stain’s cgST where re-train were performed 20 times for each cgST. Strains on the x axis were stained the same color as in Fig 2 with cgST corresponding color. Number on the x axis is the average AUROC over 20 re-trains in predicting the cgST.

**Supplementary Fig 23:**
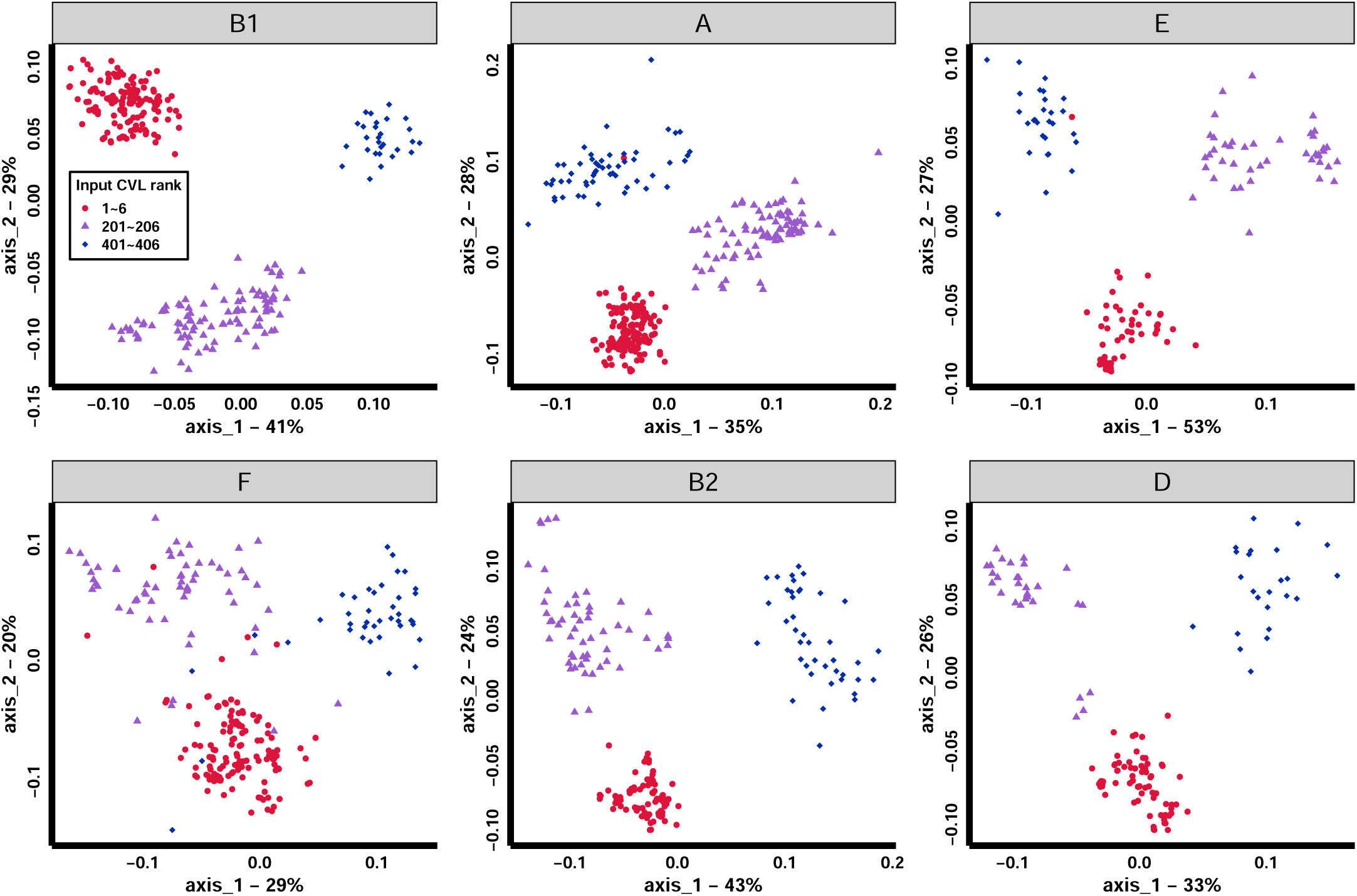
*E. coli* 3-mer difference obetween core gene combinations within each cgST. Each plot is the PCoA of 3-mer patterns of genes from CVL window of 1 to 6 (red dot), 201 to 206 (purple triangles), and 401 to 406 (blue square). All group pairwise significance test using permanova in each plot resulted in P < 0.001 significance.

**Supplementary Fig 24:**
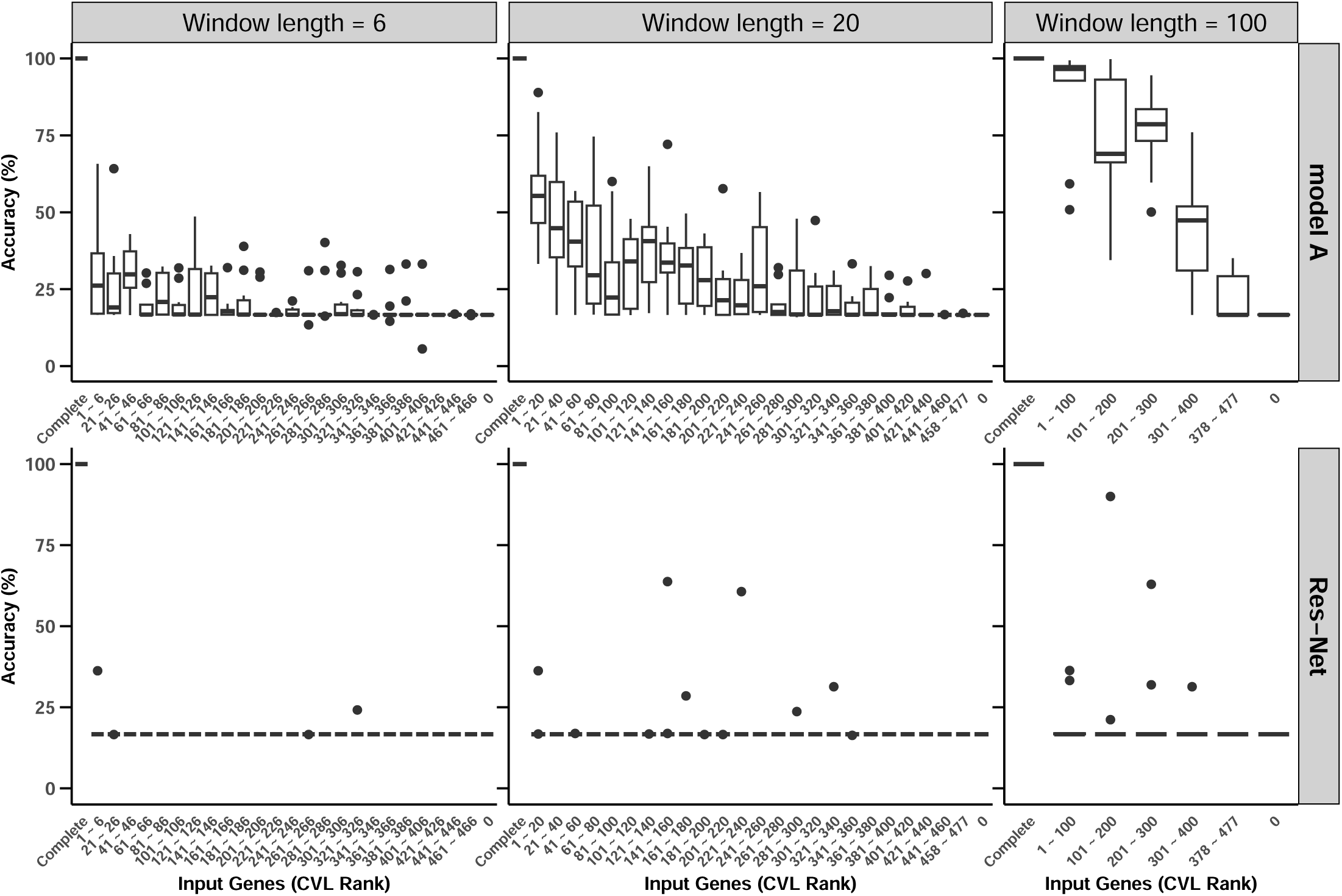
*E. coli* re-prediction performance by input of core genome subsets. two groups of models were pre-trained using complete core genome for cgST prediction, where one group used structure of model A (first row) in Supplementary Fig 8 and a Res-Net like structure (Supplementary Fig 21) for another group (second row). Ten pre-trained models from each group performed re-prediction using different subset of input. By using CVL window length of 6 (first column), 20 (second column), and 100 (thrid column) genes, re-prediction shows low accuracy while high test set performance models could be trained using same input combinations.

**Supplementary Fig 25:**
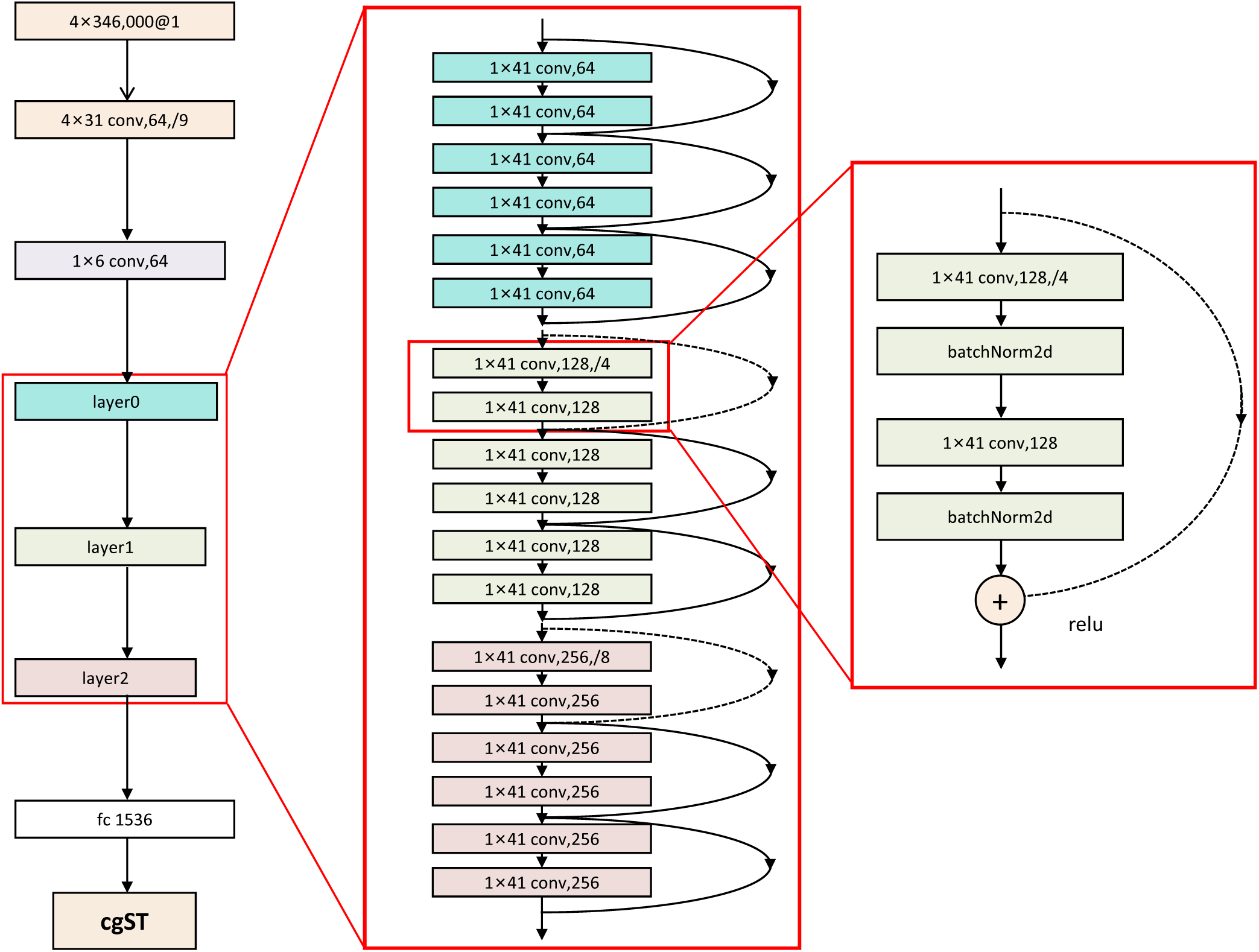
*E. coli* Res-Net like structure used for re-prediction test in Supplementary Fig 21. This customized structure input dimension is the same as complete core genome of *E. coli* Dataset II and applied 2 convolutional layer first to achieve certain dimension reduction. The model subsequently performs a Res-Net computation with 3 blocks and each block includes a cascade of 3 sections. Each section would perform residue connection, batch normalization and ReLu. Downsampling were performed between each block and a final fully connected layer helped softmax for cgST prediction.

## Notes

### Competing Interest Statement

The authors have declared no competing interest.

### Summary of Updates

We added data and in silico experiment demonstrating ultra-long distance genes-to-genes correlation to compare with gene co-occurrence and Linkage Disequilibrium.

